# GHOSTs reveal a GFP-family lineage beyond marine animals that lacks Tyr66 but preserves a buried charge network

**DOI:** 10.64898/2026.07.29.741452

**Authors:** Tanya L. Schneider, Marc Zimmer

## Abstract

Every characterized wild-type fluorescent protein (GFP family, Pfam PF01353) comes from a marine animal, and every one builds its chromophore with a central tyrosine, Tyr66 in *Aequorea victoria* GFP (avGFP). A structure-based search of the AlphaFold database for the GFP fold, followed by sequence searches, revealed a previously unrecognized lineage in bacteria, fungi, a lichen-forming fungus and corals. They carry methionine, valine, threonine or phenylalanine at the position equivalent to Tyr66. We call them GHOSTs, for GFP Homologs Omitting the Signature Tyrosine. Although they share only 23 to 35% sequence identity with avGFP over aligned residue pairs, GHOSTs retain the chromophore-site glycine and both of avGFP’s catalytic residues, the equivalents of Gly67, Arg96 and Glu222, and lack only the tyrosine. The catalytic arginine and glutamate are joined by a second arginine, a second glutamate and a tyrosine in a buried cluster we call the R2E2 core. Conventional fluorescent proteins can carry the same five residues, but in GHOST models they are packed far more closely and buried more deeply (Arg182 CZ to Glu134 CD, 4.4 versus 16.7 Å). Molecular dynamics supports a compact network that excludes water once the chromophore ring has formed, and the five core residues are among the most slowly evolving sites in the family, slower than equally buried positions. A similar geometry occurs in predicted DUF2490 barrels of a different fold. GHOSTs are the first members of the GFP family found outside marine animals, and reveal a conserved buried architecture whose biochemical role is unknown.

## Introduction

Green fluorescent protein (GFP) allows researchers to watch proteins inside living cells.^1–3^ Three consecutive residues on a central helix form a chromophore inside an eleven-stranded β-barrel^4^. In avGFP, Ser65, Tyr66 and Gly67 provide the chromophore atoms, and Arg96 and Glu222 help the maturation reaction.^5–7^ Tyr66 is especially important: its aromatic side chain becomes the light-absorbing part of the chromophore, thereby making GFP a genetically encoded tracer.

The proteins that taught us this chemistry were found in marine animals, like jellyfish, corals, copepods and lancelets.^8,9^ They were most commonly discovered by researchers looking for fluorescence, then isolating the gene. A GFP-family protein that does not produce an obvious fluorescent signal could easily escape this search. Previously reported bacterial GFP sequences have reflected contamination from laboratory vectors rather than genuine bacterial proteins.^9^

The GFP fold itself is not confined to fluorescent proteins. Nidogen contains a related β-barrel, but it lacks the chromophore-forming machinery.^9^ We asked whether a different branch might retain both the GFP barrel and the catalytic machinery while losing the canonical tyrosine. We also asked whether such proteins might occur beyond the marine animals that have dominated GFP research.

A search based on predicted structure gave us both answers. It uncovered GFP-family homologs in soil bacteria, other environmental microbes, fungi, a lichen-forming fungus and corals. We call them GHOSTs: GFP Homologs Omitting the Signature Tyrosine. Their defining feature is not simply a missing Tyr66-equivalent. It is what remains inside the barrel: a buried arrangement of two arginines, two glutamates and a tyrosine. We call this the R2E2 core (Fig. 1). Here we show how GHOSTs relate to GFP, how their core differs from that of conventional fluorescent proteins, and show that a similar chemistry-oriented geometry occurs in an unrelated fold.

**Fig. 1.**
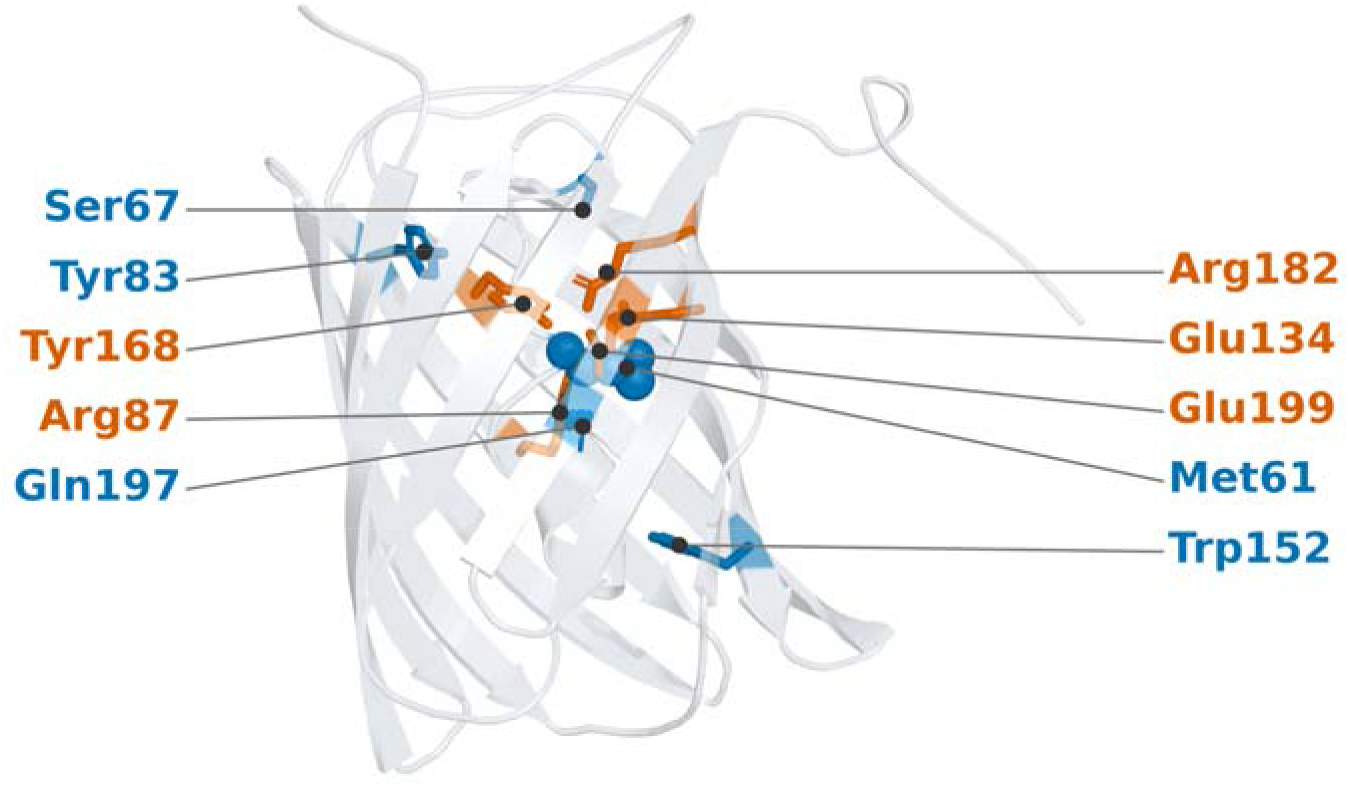
A GFP-family homolog from soil. AlphaFold model of the A0A951 β-barrel (gray cartoon). Side chains of the R2E2 core (Arg87, Glu134, Tyr168, Arg182 and Glu199) are shown as orange sticks and labeled in orange, and those of conserved Ser67, Tyr83, Trp152 and Gln197 as blue sticks labeled in blue. Fig. 3 shows the buried contacts in detail. Met61, at the position equivalent to avGFP Tyr66, is shown as blue spheres.

## Materials and Methods

### Discovery and family validation

Full protocols, software versions and sensitivity analyses are given in the Supporting Information. GFP-fold candidates were identified with Foldseek^10^ searches of the AlphaFold Protein Structure Database^11,12^, using avGFP (PDB 1EMA)^13^ and A0A951 as queries. Annotated fluorescent proteins and FPbase^14^ matches were excluded from the complete-barrel set; full-length candidates were 180–320 residues long, while three informative fragments were retained separately. BLASTp^15^ and HMMER/PyHMMER^16,17^ searches of UniProtKB^18^, MGnify^19^ and NCBI^20^ expanded the family. A 157-bit profile threshold was used to curate 22 complete barrels and three fragments (Supplementary Table 1). Equivalent residues were assigned by structural alignment with TM-align^21^ and US-align^22^, not sequence alignment. Percent identities are reported throughout over aligned residue pairs and were computed with BLOSUM62^23^ and Biopython^24^. To assess contamination, sequences were compared with FPbase using MMseqs2^25^ and screened for 15 vector motifs. The founding bacterial locus was checked for continuous read coverage, consistent gene boundaries and genomic context (Supplementary Note 1).

### Chromophore census and phylogenetics

AlphaFold hits were filtered for the GFP fold (TM-score ≥ 0.5) and the structurally mapped Gly67, Arg96 and Glu222 positions. FPbase^14^ entries were classified as wild type or engineered using curated parent relationships. Four comparison panels are used below and are not interchangeable: 194 FPbase wild-type proteins with an assignable chromophore position for the tyrosine census, 349 wild-type fluorescent proteins for the sequence logos of Fig. 2, 896 fluorescent proteins for the Pro64 enrichment test and 617 PF01353 proteins for the five-residue identity search. The primary analysis comprised 25 GHOST records and 29 comparison sequences (54 total). Sequences were aligned with MAFFT L-INS-i^26^, trimmed with trimAl^27^ (gap threshold 0.2; 228 retained columns), and analyzed in IQ-TREE^28^ under LG+G4 selected by ModelFinder^29^. Branch support was assessed with 1,000 ultrafast-bootstrap^30^ and SH-aLRT replicates. Constrained topologies were compared with approximately unbiased tests^31^ using 10,000 RELL replicates.

**Fig. 2.**
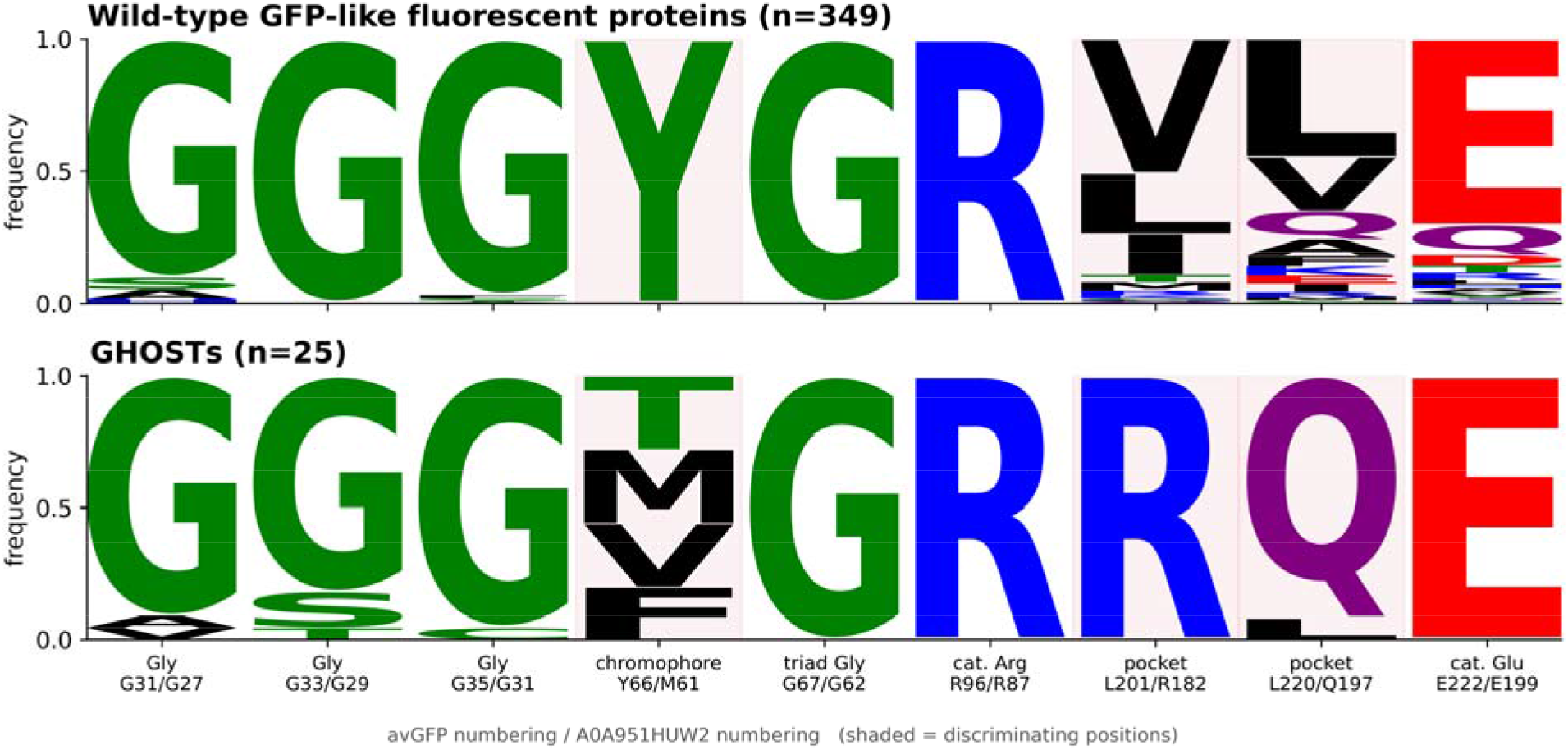
Conserved and different residues in fluorescent proteins and GHOSTs. Sequence logos compare nine positions in 349 wild-type fluorescent proteins (top) and 25 GHOST records (bottom). Letter height indicates how often each amino acid occurs. Both groups conserve the glycines important for folding and the residues involved in chromophore formation. The three shaded positions highlight key differences: fluorescent proteins contain Tyr66, Leu201 and Leu220, whereas GHOSTs have other residues at these positions. An expanded search recovered additional full-fingerprint GHOST candidates, none of which contained tyrosine at the corresponding position (Supplementary Table 8).

### Predicted structures and core geometry

Missing structures were predicted using AlphaFold2^32^ through ColabFold v1.6.1^33^. Minimum functional-heavy-atom distances defined salt bridges (<4.0 Å) and hydrogen bonds (<3.5 Å). Relative solvent accessibility was calculated with the Shrake–Rupley method^34^ in Biopython^24^, normalized to residue-specific maxima^35^. Figure 4 compares 16 GHOST models with 12 conventional fluorescent-protein models containing the same five core residues and predicted using the same pipeline; the arginine-contact and Poisson-Boltzmann comparisons use 17 GHOST models. Five independently predicted ColabFold models each of A0A951 and coral A0A7U3V473 were also examined for contact robustness (Supplementary Methods and Table 9).

**Fig. 3.**
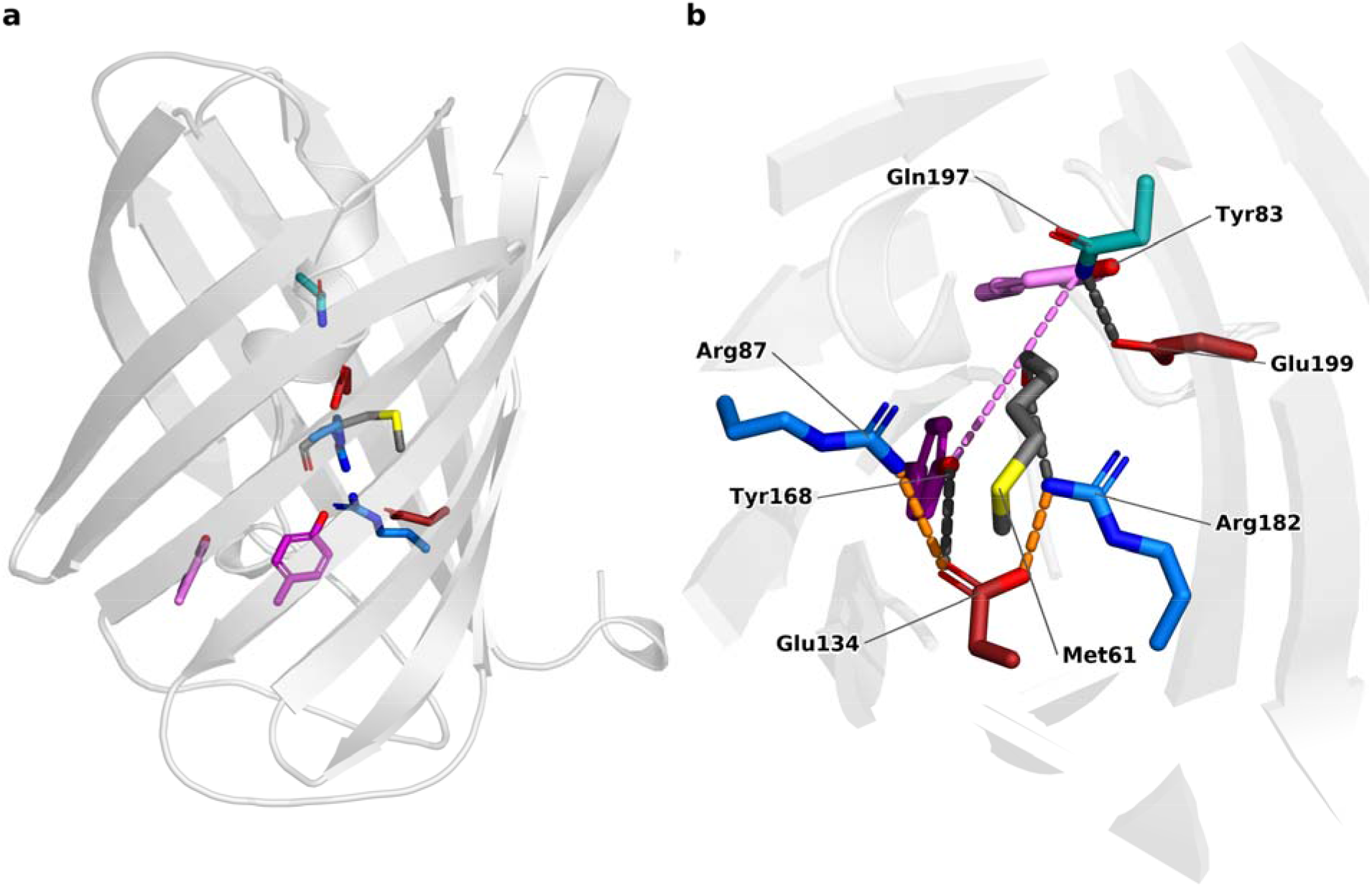
The R2E2 core of A0A951. (a) Position of the buried network within the β-barrel. (b) Close-up of the core: Arg87 and Arg182 (blue), Glu134 and Glu199 (red) and Tyr168 (dark purple), with Tyr83 (pink), Gln197 (teal) and Met61 at the Tyr66-equivalent position (gray, sulfur yellow). Orange lines denote salt bridges and dark gray lines hydrogen bonds. The purple line marks the Tyr83-Tyr168 aromatic contact.

**Fig. 4.**
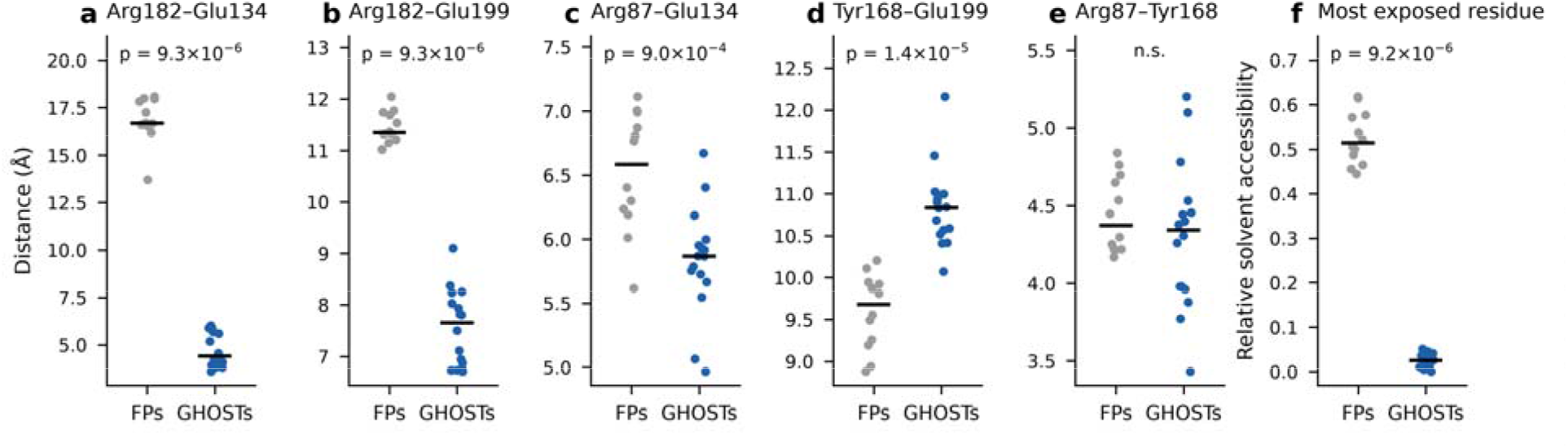
The same residues, assembled differently. Functional-atom distances (a-e; Arg CZ, Glu CD, Tyr OH) and relative solvent accessibility of the most exposed of the five core residues (f) in 16 GHOST models (blue) and 12 predicted conventional fluorescent proteins carrying the same five residues (FPs, gray). Each point is one structure, bars are medians, and *p* values are from two-sided Mann-Whitney U tests. In GHOSTs Arg182 lies close to both Glu134 (a) and Glu199 (b), Arg87 lies closer to Glu134 (c), and all five residues are buried (f). Tyr168-Glu199 is the exception and is longer in GHOSTs (d); Arg87-Tyr168 does not differ (e). The distinguishing feature is therefore the arrangement and burial of the network, not its amino-acid composition. All comparisons use predicted structures.

### Molecular dynamics and electrostatics

Explicit-solvent simulations used OpenMM^36^, Amber ff14SB^37^, TIP3P water^38^, 0.15 M NaCl, 300 K, 1 bar and a 2 fs timestep. Nonstandard chromophore species used GAFF2 parameters, an updated version of the general Amber force field,^39^ with AM1-BCC^40^ charges, both from AmberTools^41^. Three independent 100 ns replicas were run for each system. Four A0A951 systems were simulated: a cyclized immature-ring model and the linear pre-cyclization protein, each with Glu199 charged and with Glu199 protonated. The coral A0A7U3V473 F67Y model and DUF2490 protein A0A8J3CUU4 were simulated in the same way.

Contacts, hydration and fluctuations were analyzed with MDTraj^42^. PROPKA^43^ was used to predict p*K*_a_ shifts, and PDB2PQR/APBS^44^ to perform two Poisson–Boltzmann analyses at pH 7: an AMBER-parameterized single-structure analysis of the A0A951 network, and a PARSE-parameterized comparison of chromophore-site carbonyl potentials in 17 GHOST, 18 general fluorescent-protein and 12 core-matched fluorescent-protein models, with chromophore-triad charges zeroed.

### Evolutionary constraint

Site-specific rates were estimated in IQ-TREE^28^ under LG with 20 discrete Gamma rate categories using the 25-sequence MAFFT alignment (325 columns) and its maximum-likelihood tree. The 218 columns mapping to A0A951 were analyzed. Rates at the five core positions were compared with the remaining 213 positions and with equally buried sites at relative accessibility thresholds of 0.01–0.20. One-sided Mann–Whitney U tests and 100,000 five-site permutations were used; ConSurf^45^ provided an independent conservation estimate.

### Cross-fold motif and genomic-context searches

The five core residues were searched against other folds using RCSB Structure Motif Search^46^ and FoldDisco^47^. Of 4,234 DUF2490 (PF10677) members, the 2,789 with AlphaFold models were screened for a functional-atom RMSD ≤1.8 Å, two salt bridges <4.0 Å and aromatic engagement <4.0 Å. Signal peptides were assessed with SignalP 6.0^48^. The founding contig and 11 additional loci were annotated with the NCBI pipeline^49^ and Prodigal/Pyrodigal^50,51^. BWA-MEM^52^ and SAMtools^53^ were used for read mapping; geNomad^54^, CheckV^55^, VirSorter2^56^, DefenseFinder^57^ and PADLOC^58^ were used to screen for mobile elements and defense systems (Supplementary Note 1).

### Statistical analysis

Tests were two-sided unless stated otherwise. The five core residues were selected from the structural analysis before the evolutionary-rate comparison, but the test was not preregistered. Analyses used SciPy^59^, NumPy^60^ and Biopython^24^; further parameters and sensitivity analyses are provided in the Supporting Information.

## Results

### A GFP-family protein turns up in soil

We searched predicted protein structures with avGFP (PDB 1EMA) using Foldseek.^10,13,61^ After excluding annotated fluorescent proteins, we found A0A951HUW2 (A0A951), a 218-residue protein from an alphaproteobacterial metagenome-assembled genome collected in Brazilian rhizosphere soil.^62^ The predicted structure overlays avGFP with a TM-score of 0.844 and an RMSD of 2.09 Å over 202 residues. Their sequences share only 23% identity over aligned residue pairs (Supplementary Table 11). Nevertheless, the central chromophore forming machinery is recognizable. A0A951 Gly62 aligns with avGFP Gly67, Arg87 with Arg96, and Glu199 with Glu222.^63^ The characteristic scaffold glycines are also retained.^64^ But the residue equivalent to Tyr66 is Met61 (Fig. 1). The protein keeps much of the apparatus associated with chromophore formation while changing its most familiar component.

A soil metagenome could contain an assembly error or a stray laboratory GFP sequence. We therefore traced the gene back to its source. Two gene callers assign the same open reading frame. Read coverage is continuous, paired reads span both gene boundaries, and the surrounding contig has bacterial gene content without eukaryotic markers. The evidence supports a genuine bacterial locus rather than a misassigned GFP construct (Supplementary Note 1).

### GHOSTs extend the GFP family beyond marine animals

Searches seeded with A0A951 recovered 25 GHOST records: 22 complete barrels and three partial fragments. All 25 cover the chromophore-equivalent site, but only the 22 complete proteins allow every network position to be scored. Among those 22 are 14 bacterial proteins, five fungal proteins, two coral proteins and one reef metagenomic record. The fungal set includes a sequence attributed to a lichen-forming fungus. Five records are marine. The other 17 include soil, groundwater and hot-spring samples, although habitat information is incomplete for some entries (Supplementary Table 1). Among the 12 records with a recorded geographic origin, at least one came from each of the seven continents.

These are, to our knowledge, the first well-supported bacterial and fungal members of the GFP family that retain its characteristic central machinery. They are not merely unrelated proteins with a barrel of similar shape. A0A951 has almost the same structural similarity to conventional fluorescent proteins (TM-score 0.862) as it has to other GHOSTs (0.870). A 54-sequence maximum-likelihood analysis places all 25 GHOST records in one clade, with SH-aLRT support of 92.9 and ultrafast bootstrap support of 98 (Supplementary Fig. 1). A constrained search returns the same monophyletic group.

The relationships within that group are less secure. The fungal sequences cluster, but their placement relative to bacterial sequences depends on taxon sampling. A test forcing the sampled prokaryotic sequences into one group is rejected in our analysis (approximately unbiased *p* = 1.1 × 10^-4^). We therefore do not infer a single transfer route or the direction of transfer from this tree alone. An independent assembly-wide genomic census, posted on 3 November 2026, also recovers the family and greatly extends its taxonomic range.^65^ It does not recover an exclusive prokaryotic clade, although its test does not significantly reject one. The datasets agree on the family’s existence but are less decisive about its deeper history.

### The canonical Tyr66 is absent, but its partners remain

Every wild-type fluorescent protein builds its chromophore around a tyrosine. In FPbase, every curated wild-type fluorescent protein with an assignable chromophore position carries tyrosine there (n = 194, after excluding channelrhodopsin 2, a retinal-binding opsin that the structural assignment matches spuriously rather than a GFP-fold protein). Every other residue found at this position was introduced in the laboratory. Fluorescent proteins built on other scaffolds, among them the bilirubin-binding eel protein UnaG and the phytochrome- and flavin-derived reporters, fall outside Pfam PF01353 and outside this comparison.

The pattern holds in the AlphaFold database. Of 6,533 structural matches to avGFP, 2,682 have a genuine GFP fold (TM-score ≥ 0.5), but most of these lack the chromophore-forming machinery. Requiring the Gly67/Arg96/Glu222-equivalent trio leaves 457 proteins, and 446 of them (97.6%) carry tyrosine. None of the 11 exceptions is a natural fluorescent protein. Three are laboratory mutants deposited under their source organisms. Two are uncharacterized GFP-like proteins from the lancelet *Branchiostoma belcheri* (UniProt A0A6P5AE48 and A0A6P4ZFR3) with phenylalanine at this position, neither known to fluoresce. The remaining six are GHOSTs. Across the complete GHOST barrels the Tyr66-equivalent position is Met, Val, Thr or Phe, and none has tyrosine.

The surrounding window is also distinctive. In A0A951 it is Met61-Gly62-Xaa63-Pro64. Gly62, corresponding to the canonical Gly67, is present in all 25 records. Pro64, three positions after Met61, is also present in all 25, compared with 16 of 896 proteins in the fluorescent-protein panel (Fisher exact *p* = 1.7 × 10^-38^). The significance of this enrichment must be interpreted with care because related sequences are not independent evolutionary samples. Nevertheless, the sequence pattern is striking. GHOSTs do not simply lack one residue from an otherwise ordinary fluorescent-protein motif (Figs. 2 and 3).

### A conserved five-residue network occupies the barrel interior

GHOSTs preserve much more than the canonical catalytic pair. In the A0A951 model, Arg87, Glu134, Tyr168, Arg182 and Glu199 form a compact, buried arrangement that we call the R2E2 core (Fig. 3). All five residues are invariant across the 22 complete GHOST proteins. They are also retained in the additional full-length candidates examined during the expanded search. Ser67 and Trp152 are conserved as well, and Gln197 is present in 20 of the 22 original complete members.

The arrangement connects the original GFP catalytic residues to additional charged and aromatic groups. In A0A951, the closest heavy atoms of Glu134 and Arg182 lie 2.60 Å apart, and those of Glu134 and Arg87 3.26 Å. Arg182 is 2.81 Å from the Met61 backbone carbonyl, while Gln197 contacts Glu199 at 2.69 Å. Tyr168 sits within this buried environment. The core residues are almost completely buried, with solvent-accessible surface areas of just 0–8 Å^2^.

The arrangement is not confined to one predicted structure. Five replicate ColabFold predictions for A0A951 and a coral GHOST, with side chains relaxed, retain the principal contacts. The weaker Arg87-Glu134 interaction varies near the salt-bridge cutoff. Local prediction confidence is high (pLDDT 95 to 99). These comparisons support a robust structural hypothesis: GHOSTs conserve a specific buried architecture, not just five amino-acid identities.

### The difference between GHOSTs and fluorescent proteins is the packing, not the ingredients

The five R2E2 residues are not unique to GHOSTs. We found the same residue identities in 50 of 617 PF01353 proteins, including 45 conventional tyrosine-bearing fluorescent proteins. This is an important control. GHOSTs are distinguished by a broader fingerprint that includes Ser67, Trp152 and the non-tyrosine chromophore site. The shared residues are not arranged in the same way in the two groups.

We compared 16 GHOST models with 12 predicted fluorescent-protein structures that carry all five core residues and could be mapped unambiguously. In GHOSTs, the core forms a more connected, buried assembly (Fig. 4). The median Arg182-Glu134 functional-atom distance, measured from Arg CZ to Glu CD as in Fig. 4, is 4.40 Å, compared with 16.67 Å in the matched fluorescent proteins (*p* = 1 × 10^-5^). Arg182-Glu199 is also closer: 7.65 Å versus 11.34 Å. Mean relative solvent accessibility across the five positions is 0.01 in GHOSTs and 0.11 in the controls. At least one position is substantially exposed in many of the conventional proteins. Neither group is a set of independent evolutionary samples, so we repeated every comparison after collapsing each group to sequence clusters. The separation persists at every identity threshold down to 50%, where ten family clusters and three control clusters remain completely separated on all five measurements (Supplementary Table 12).

Not every distance is shorter in GHOSTs. Arg87-Tyr168 is nearly unchanged (4.34 Å versus 4.37 Å), and Tyr168-Glu199 is farther apart (10.83 Å versus 9.68 Å). Not all five side chains collapse into the smallest possible pocket. The distinctive feature is a particular packing pattern that links the two arginines to the glutamate environment and keeps most of the assembly buried.

The chromophore-site backbone also lies closer to an arginine in GHOST models. The nearest guanidinium nitrogen is a median of 3.22 Å from the corresponding carbonyl in 17 GHOST structures. The distance is 3.58 Å in 12 composition-matched fluorescent-protein models and 4.12 Å in a wider control panel. All comparisons used the same structure-prediction pipeline. The corresponding atom in a mature fluorescent-protein crystal structure belongs to a different chemical group, so it cannot be used as a direct geometric control.

### The network persists in simulations and largely excludes water

Molecular dynamics tested whether the predicted contacts survive thermal motion. We ran three independent 100-ns explicit-solvent replicas of the cyclized A0A951 immature-ring model with Glu199 charged and three with Glu199 protonated, and the same two sets for the linear pre-cyclization protein. The contact measurements below are from the cyclized model. The Arg182-Glu134 and Arg87-Glu134 contacts persisted in essentially every analyzed frame. So did the Arg182 contact to the Met61 carbonyl. The Gln197-Glu199 interaction remained present in 97 to 100% of frames. Core side-chain fluctuations were small (0.44 to 0.77 Å). The Arg182–Glu134-equivalent contact also persisted in simulations of an F67Y coral GHOST model with a mature chromophore.

The center of the network was water-excluded once the chromophore ring had formed, and not before. In the cyclized model, with Glu199 either charged or protonated, no simulated water molecule bridged either of the two principal cation-anion contacts, Arg87, Arg182 and Glu134 contacted water within the defined hydrogen-bond cutoff in at most 2% of frames, and the Met61 carbonyl was not hydrated by this measure. The linear pre-cyclization protein behaved differently: the same three residues were hydrated in 16 to 29% of frames, the Met61 carbonyl in 6 to 20%, and a bridging water was present at each of the two salt bridges in roughly one frame in six. Glu199 remained accessible to water at the edge of the network in three of the four systems. In these simulations the core is dry in the cyclized state and hydrated in the linear one, so the dryness is a property of the modeled cyclized state rather than a constitutive feature of the fold. These results support a compact, partly insulated electrostatic environment under the modeled conditions. They do not by themselves establish a chemical reaction (Supplementary Note 5).

### Evolution preserves the network more strongly than the chromophore position

The five core positions are invariant in every complete member of the original GHOST set. Their estimated evolutionary rates place them among the slowest-changing sites in the 218-residue barrel (Fig. 5). Under the 20-category rate model, their rates range from 0.085 to 0.099, with percentiles of 2 to 4. The median rate is 0.088 for the five core sites and 0.732 for the other 213 positions (one-sided Mann-Whitney *p* = 9.6 × 10^-5^).

**Fig. 5.**
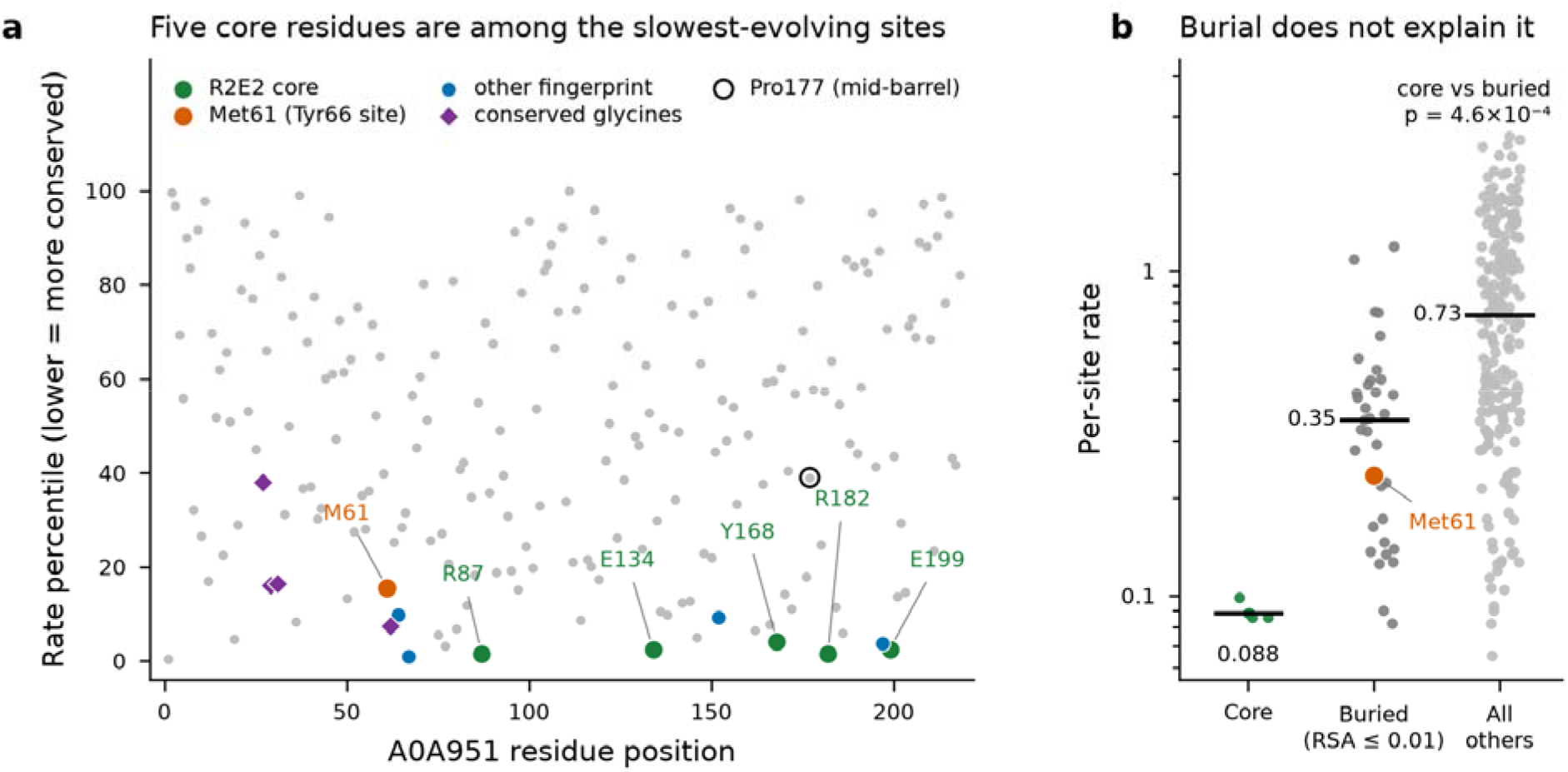
Evolutionary constraint concentrates on the R2E2 core. (a) Empirical-Bayes per-site evolutionary-rate percentiles for the 218 A0A951-mapped barrel positions, from the 25-record family alignment under LG with a 20-category discrete Gamma model. Lower percentiles indicate slower evolution. The R2E2 core (green) falls at the 2nd to 4th percentiles and Met61, the Tyr66-equivalent site (orange), at the 16th. Other fingerprint residues (Pro64, Ser67, Trp152, Gln197), the conserved glycines (Gly27, Gly29, Gly31, Gly62) and a mid-barrel control (Pro177) are shown for comparison. (b) Per-site rates for the five core positions, for the 35 other positions as buried as the core (relative solvent accessibility 0.01 or less in the A0A951 model) and for all other positions. Bars are medians. Met61, itself fully buried, is marked. The comparison is descriptive of evolutionary constraint, not proof of catalytic function.

The Tyr66-equivalent position is more variable. In A0A951, Met61 has an estimated rate of 0.235 and lies at the 16th percentile. Thus, the family tolerates several residues at the canonical chromophore site while retaining the same buried core. This contrast is consistent with strong structural or functional constraint on the network. It does not, on its own, identify the function. Buried residues often evolve slowly for reasons of stability, so we repeated the test against positions as buried as the core. Burial accounts for part of the effect but not most of it. The 35 other positions with relative solvent accessibility of 0.01 or less have a median rate of 0.348, four times that of the core, and the core remains more constrained than them (one-sided Mann-Whitney *p* = 4.6 × 10^-4^; permutation *p* ≤ 10^-5^). The result holds at every burial threshold tested. Only three buried positions evolve as slowly as the core: Ser67, Phe77 and Gln197, all neighbors of the network. Met61 is fully buried too, yet ranks only at the 32nd percentile among buried sites (Fig. 5b).

### The local electrostatics are distinctive, but not diagnostic by themselves

Poisson-Boltzmann calculations predicted a positive potential at the chromophore-site carbonyl. The mean was 36.2 kT/e in 17 GHOST models and 19.4 kT/e in 18 general fluorescent-protein models (*p* = 3.0 × 10^-4^). Removing the arginine charges reversed the sign in both groups. But most members of that general control set did not carry the second arginine. When we instead used 12 fluorescent-protein models with the matched five-residue core, the comparison was 36.2 versus 24.8 kT/e and was not statistically significant (*p* = 0.06). The electrostatic potential therefore does not independently define GHOSTs. The more convincing distinction is the geometry and burial of the network.

PROPKA predicts an upward shift in the p*K*_a_ of a buried arginine in several GHOST models, consistent with a strongly perturbed electrostatic environment. These are model-based predictions, not measured protonation equilibria. Taken together, the geometry, burial, molecular dynamics and electrostatics suggest a specialized internal environment. They do not yet show that it transfers electrons or protons.

### The fold tolerates different residues at the missing Tyr66 site

The chromophore-equivalent position varies naturally in GHOSTs, so we tested whether its identity changes the predicted fold. ColabFold models of A0A951 and five residue-61 variants (Val, Phe, Tyr, Thr and His) retained closed GFP-like barrels across 96 models, with and without GFP templates. In particular, restoring tyrosine (M61Y) caused no reproducible pocket expansion or close clash in the models. This suggests that the barrel can accommodate a tyrosine at that position. It does not establish that M61Y forms a chromophore or that the natural proteins are nonfluorescent (Supplementary Note 8).

### An unrelated protein family has a similar buried network

If this geometry is useful, it might recur outside the GFP fold. We searched for the five-residue motif without requiring a GFP-like structure. Searches of experimental PDB entries and approximately 1.06 million RCSB computed models did not recover a convincing independent buried match under our filters. A much larger FoldDisco search of approximately 53 million AlphaFold cluster representatives did. It identified the GHOST family, then a second group of proteins belonging to DUF2490 (PF10677).^46,47^

DUF2490 proteins are predicted outer-membrane β-barrels. They lack GFP’s central helix and are not GHOSTs. Among 2,789 DUF2490 members with AlphaFold models, 698 carried the required residue classes in approximately the same constellation. Of these, 238 met the more stringent geometric definition of an engaged network, with two salt bridges and a nearby aromatic group. In representative models, the five functional groups superpose on the GHOST core with an RMSD of 1.2 to 1.35 Å, even though the overall folds align poorly (TM-score about 0.37). A separate barrel architecture therefore appears able to organize a similar set of buried functional groups.

The similarity has limits. In molecular-dynamics simulations of one DUF2490 member, one arginine-glutamate salt bridge persisted, but contacts involving the second arginine varied greatly between replicas. The aromatic side chains were also much more mobile than in A0A951. Thus, DUF2490 provides evidence for recurrence of the predicted geometry, not proof of an identical, rigid arrangement or a common function. No experimental DUF2490 structure or biochemical role for the motif is yet available.

## Discussion

GFP gives the crystal jellyfish much of its green light. Its use as a protein tracer helped researchers discover a remarkable family of fluorescent and colored proteins in marine animals. It also defined what researchers expected a GFP-family protein to look like. GHOSTs challenge that expectation. They retain the GFP barrel and the residues associated with chromophore maturation, yet they lack the canonical Tyr66-equivalent. Their genes also occur in soil bacteria, other environmental microbes and fungi. The GFP family is not confined to the organisms in which fluorescent proteins are found.

The family relationship is supported by more than a barrel-shaped prediction. GHOSTs share GFP’s distinctive scaffold, central glycine and catalytic Arg/Glu positions. Structural comparisons place them close to conventional fluorescent proteins, and sequence-based phylogenetic analysis recovers a supported GHOST clade. The founding bacterial locus also survives a direct provenance check. These independent observations support a genuine, previously overlooked GFP-family lineage.

The more surprising finding is what GHOSTs conserve. Their Tyr66-equivalent position varies among several non-tyrosine residues. Yet Arg87, Glu134, Tyr168, Arg182 and Glu199 are invariant across the complete family examined. In predicted structures they form a compact, buried network. Molecular dynamics preserves much of that geometry and keeps water away from its center. The same residues can occur in ordinary fluorescent proteins, but they are not packed or buried in the same way. Thus, the family is defined by both its sequence fingerprint and its unusual internal architecture.

That architecture looks chemically interesting. It places a buried tyrosine in an arginine/glutamate-rich environment close to the chromophore-equivalent backbone. Such an environment could support proton or charge-transfer chemistry. But the present calculations do not establish a transfer reaction, a redox potential or a biological substrate. Even the electrostatic potential, once compared with composition-matched fluorescent proteins, is not uniquely elevated in GHOSTs.

DUF2490’s predicted outer-membrane barrels are unrelated to GFP, yet some organize a similar set of buried charged and aromatic groups. This recurrence is consistent with a useful physical arrangement. It is not evidence that the two families share the same activity. In fact, the DUF2490 network is more mobile in our simulations. Testing whether these two folds exploit similar electrostatic properties would be an informative next step.

## Conclusions

GHOSTs are a GFP-family branch that fluorescence-guided discovery overlooked. They preserve a buried architecture that remains conspicuous across divergent sequences and appears again in an unrelated fold. The GFP barrel may have more chemical possibilities than its best-known chromophore suggests.

## Supporting Information

Supplementary Methods, expanded phylogenetic and structural analyses, additional figures and tables, and a guide to the supporting data and code are supplied in a separate Supporting Information file.

## Data availability

Data and code availability. The canonical family roster, with live primary accessions, UniParc identifiers for the three entries no longer in UniProtKB (A0A951HUW2, A0A1M3T1J2 and A0A931S450), taxon, completeness, fingerprint state and the analyses each record enters, is provided as Supplementary Table 1. Per-structure measurements behind Fig. 4 (functional-atom distances and relative solvent accessibility for 16 family and 12 identity-matched control models) and the per-structure Poisson-Boltzmann potentials for both control sets are provided as Supplementary Tables 9 and 10. The 54-sequence alignment, the maximum-likelihood tree with SH-aLRT and ultrafast bootstrap support, and the constraint trees and approximately unbiased test outputs are deposited with the manuscript. AlphaFold models for control proteins were retrieved from the AlphaFold Protein Structure Database (version 6). All processed data, analysis scripts and the residue-61 panel are archived at Zenodo (DOI 10.5281/zenodo.24164887), paths below and in the Supplementary Information are relative to the root of that archive.

Query and models: 1EMA; AlphaFold A0A951HUW2. Source data: BioProject PRJNA687273, MAG BioSample SAMN17140656, source-metagenome BioSample SAMN10290267, SRA run SRR8113284, contig JAFAZW010000077.1, protein MBV9931914. Genomic-context and defense-system outputs are in 09_genomic_context_defense/ (A0A951HUW2_gene_neighborhood.md, A0A951HUW2_crossmember_neighborhoods.md, A0A951HUW2_defense_systems.md and A0A951HUW2_padloc_systems.md). The family profile HMM (a0a_subfamily.hmm), its seed alignment (a0a_subfamily_seed.afa), calibration, the UniProtKB and MGnify sweep and verification of cnidarian, fungal and metagenomic hits are in 03_sequence_homology_hmm/. Structural models and the per-member master table (01_query_and_models/), structure searches, phylogenies and ancestral reconstructions, site rates, core geometry, molecular-dynamics occupancy tables (07_molecular_dynamics/), electrostatics and figures occupy folders 01 to 10. Motif searches, the DUF2490 census, fold-classification tests and SignalP outputs are in 11_motif_search_convergence/. The residue-61 mutational panel, with exact sequences, run configurations, manifests, all 96 model-level measurements, construct summaries and the analysis code, is in 12_residue61_panel/. Analysis and figure scripts are in scripts/, mapped to analyses in Supplementary Table 3. Quantum-chemical analyses are detailed in the companion study.

## Accession Codes

The founding bacterial GHOST is represented by NCBI protein MBV9931914 (original UniProtKB accession A0A951HUW2; UniParc UPI001DADDAB1). Accession identifiers for the other GHOSTs and the structures used in the analyses are provided in Supplementary Tables 1, 9 and 10.

## Acknowledgments

This work was supported in part by an Anthropic AI for Science award to M.Z., which provided the computational credits used for the structural, phylogenetic, molecular-dynamics and electrostatics analyses reported here.

## Author Contributions

M.Z. designed the study, performed the computational analyses and wrote the manuscript. T.L.S. contributed to the planning of experimental studies, interpretation of the findings, and revision of the manuscript. Both authors reviewed and approved the final version.

## Notes

The authors declare no competing financial interest.

