## Supplementary Material for "GHOSTs reveal a GFP-family lineage beyond marine animals that lacks Tyr66 but preserves a buried charge network"

### **Supporting Information**

#### **Supplementary Methods**

##### **Structural similarity search and candidate selection**

Candidate GFP-fold barrels were identified by structural-similarity search with Foldseek^10^, using the avGFP crystal structure (PDB ID 1EMA^13^) as the query against the AlphaFold Protein Structure Database^11,66^ cluster representatives^12^. A second A0A951-seeded Foldseek search used the A0A951 AlphaFold model (AF-A0A951HUW2-F1-model_v6) against the Foldseek AlphaFold/UniProt50 cluster-representative database (AlphaFold/UniProt50 cluster representatives, afdb50 version 6) in 3Di/AA mode through the public Foldseek server, accessed July 12, 2026. The downloaded Foldseek tables did not retain a separate server-build string beyond the database labels and job identifiers; searches used the public-server 3Di/AA defaults for sensitivity, prefiltering, E-value reporting and score reporting, and no additional server-side minimum E-value, coverage or score cutoff was applied before export of the ranked hits. Instead, E-values and alignment scores were retained for review, and candidate selection used scripted post hoc filters. Hits were filtered to remove the A0A951 self-hit, proteins with direct Pfam PF0135323 annotation, and proteins with fluorescent-protein-like names or FPbase^14^ matches; the latter filter was therefore conservative for name-recognizable fluorescent proteins. Full-barrel candidates were retained when the target length was between 180-320 amino acids, the aligned query span covered at least 180 of the 218 modeled query residues, and the alignment included the A0A951 barrel core around residues 55-205. The 180-320-residue length window was chosen to retain single-domain GFP-fold barrels with modest terminal extensions while excluding short fragments and obvious multidomain proteins. Fragments were rejected unless they covered the barrel core and were used only as sensitivity candidates; multidomain proteins were evaluated by the aligned barrel segment, with terminal extensions or extra domains recorded as candidate-quality flags. Initial filtering was automated from the Foldseek^10^, UniProt/Pfam and name-matching fields, followed by manual review of taxonomy, annotation, barrel coverage, duplicate status and retention of the GHOST fingerprint. Retained candidates were superposed on PDB ID 1EMA and their residues mapped to the avGFP scheme with TM-align^21^ and US-align^22^; the local US-align executable was version 20260527. The chromophore triad, the catalytic Arg96 and Glu222, and the buried network positions were assigned from these structural superpositions rather than from sequence alignment. Structural positions use the avGFP numbering (Ser65-Tyr66-Gly67, Arg96, Glu222) throughout, and A0A951 positions use its model numbering (Met61, Arg87, Glu199, Arg182, Gln197).

##### **Sequence homology search and family assembly**

Homologs were identified by BLASTp^15^ searches of the NCBI non-redundant protein database^20^ and by iterative jackhmmer and profile-HMM searches. The initial remote BLASTp search used A0A951 as the query with an E-value cutoff of 1e-3, SEG filtering enabled, max_target_seqs set to 500, and tabular output reporting accession, title, percent identity, aligned length, query and subject length, query coverage, E-value and bit score; the exact BLAST+ client version and nr access date were not retained in the analysis directory. The jackhmmer searches requested five iterations; the Swiss-Prot search converged after three iterations, and the local top-30 BLAST-hit search converged after two iterations. To define the family boundary quantitatively, we used HMMER^16^/PyHMMER^17^ to construct profile hidden Markov models from a clean ten-member GHOST bacterial seed set (A0A951HUW2, MBV9931914, *Mucilaginibacter*, Bacteroidota, *Winogradskyella*, Gaiellaceae, Thermoleophilia, Actinomycetota, Methylococcales and Opitutales representatives), with an *Acropora*-Phe sensitivity seed analyzed separately. The calibrated profile was then searched against UniProtKB^18^, Swiss-Prot, MGnify^19^/Foldseek structural-hit sets, NCBI annotation-derived candidate pools, and local curated bacterial candidates. Conventional fluorescent proteins were excluded, and each recovered cnidarian, fungal, and metagenomic candidate was individually evaluated by sequence similarity, predicted structure, and the residue present at the chromophore-defining position (Supplementary Note 2).

To search the space excluded by the structure-first step, all 689 UniProtKB entries carrying Pfam PF01353 were retrieved, the 25 roster sequences aligned with MAFFT and built into a profile with hmmbuild, and the profile searched against that set with hmmsearch (--max, E < 1e-5). Hits were aligned back to the profile with hmmalign and the A0A951 fingerprint positions read directly from the resulting match columns, which avoids the pairwise-alignment failure described above. Residue identities were scored against the five buried core positions and the eight-residue fingerprint. Recovery of the four roster members present in this set served as the positive control.

##### **Provenance and contamination screening**

Every non-reference taxon was screened against FPbase reference sequences for full-length identity, and each member was checked for FPbase membership, PDB cross-references and UniProt protein-existence level. Cross-domain members were further supported by their source records (coral transcript evidence and intron-containing fungal gene models; Supplementary Table 2). Three EGFP-carrying reference assemblies were identified as synthetic-construct contamination and excluded. The screen was subsequently rebuilt as an automated two-signal procedure and applied to the full unannotated GFP-fold cohort: MMseqs2^25^ identity against all 992 FPbase reference sequences, computed over the shorter sequence, combined with a scan for fifteen expression-vector motifs (polyhistidine, pQE MRGS-His, TEV, thrombin, enterokinase, FLAG, Strep-II, S-, HA-, Myc- and T7 tags, Gly-Ser linkers, AviTag and a SUMO C-terminus), with host taxonomy resolved through the UniProt taxonomy service rather than by genus keyword. The procedure recovers all four previously documented contaminants in this cohort and separates transgenes residing in a foreign host from engineered constructs deposited under their source organism: it flags nine of the former, five of which were not previously recorded, and three of the latter, while 874 of 886 unannotated barrels pass. No GHOST member is flagged by either signal. For the founding contig, the audit also recorded coding density, the upstream ribosome-binding motif, agreement of gene boundaries and start sites, boundary-spanning read pairs, coverage discontinuities and soft clips, and the probability outputs of geNomad together with the VirSorter2 and CheckV calls. Re-checking the roster against current UniProtKB showed that four of the ten UniProt-derived records had been deleted as 'not part of a reference proteome' rather than merged into another accession: A0A951HUW2 (the founding sequence), A0A931S450, A0A146F2F0 and A0A1M3T1J2. Each sequence is intact and retains an active primary record that is cited in its place (MBV9931914, MBI2083722, GAT20325 and OJZ80615; UniParc UPI001DADDAB1, UPI001DD0F21D, UPI0007947AEC and UPI000924AB72). A0A146F2F0 (282 residues) and A0A7R7WR58 (254 residues) are the same *Aspergillus kawachii* protein under two gene-model start calls, the shorter contained exactly within the longer from offset 28. This is the only redundant pair among the records and both carry an identical fingerprint, so A0A146F2F0 was withdrawn without changing any invariance count. A0ABR1SEL9, which carries the complete eight-position fingerprint with valine at the chromophore position, was added as a separate member. The roster therefore comprises 22 complete barrels and three fragments.

For the placement test of the cephalochordate phenylalanine locus, the 48 unique sequences of the taxon-targeted alignment were degapped, combined with A0A6P5AE48 and the tyrosine-bearing control C3YKQ3, realigned with MAFFT in --auto mode, and an approximate-maximum-likelihood tree inferred with FastTree under the LG model with Shimodaira-Hasegawa-like local support. This is a placement check and not a re-inference of Supplementary Fig. 4: it uses a different alignment and a different inference program from the maximum-likelihood analysis reported there, and it does not provide ancestral-state reconstruction. The reported conclusion is limited to which clade the locus joins.

Sequences were aligned with FAMSA^67^ through PyFAMSA 0.7.0, using duplicate retention where duplicate labels or sequences were required for taxon auditing. No manual alignment editing was performed. For the cross-domain phylogeny shown in Supplementary Fig. 4, sequences were projected onto the A0A951-aligned columns used in the ConSurf/broad-context alignment, yielding a 49-sequence amino-acid alignment of 218 retained sites. For the expanded AFDB50-informed tree, the untrimmed alignment contained 508 columns and was reduced with a scripted 50% occupancy filter to 227 retained sites. Partial source sequences were retained only when they covered the GFP barrel core; residues outside their aligned span were treated as gaps.

Residue-position equivalences to avGFP used throughout this work were assigned from structural superposition rather than from sequence alignment. That choice is load-bearing at these identities. For one family member, three alignment routes disagreed on the identity of the Glu222-equivalent position, returning glutamate, valine and leucine respectively, whereas superposition-free internal distances measured from the chromophore within each structure placed a glutamate carboxylate 4.77 A from the chromophore ring centroid against 4.68 A in the avGFP crystal. Global sequence alignment is therefore unreliable for the carboxy-terminal catalytic positions at the identities found here, and all such assignments reported here were confirmed structurally.

##### **Additional phylogenetic sensitivity analyses**

Historical sensitivity analysis, not the primary 54-sequence tree: Maximum-likelihood trees were inferred in IQ-TREE 3.1.334, with the best-fit substitution model selected by ModelFinder^29^, branch support from 1000 ultrafast-bootstrap replicates^30^, and 1000 SH-aLRT replicates. For the cross-domain phylogeny shown in Supplementary Fig. 4, ModelFinder selected LG+I+G4 by BIC for the 49-sequence, 227-site alignment. The earlier record of this run as VT+I+G4 over 218 sites was an error; the archived inference is the 49-sequence, 227-site run under LG+I+G4, and re-inference of that alignment reproduces its topology exactly, at zero Robinson-Foulds distance across all 45 splits. The complete branch-length phylogram of this historical run is provided as Supplementary Fig. 4. The expanded AFDB50-informed tree contained 68 sequences and 227 retained sites, and ModelFinder selected Q.PFAM^68^+F+R4. Primary ML trees were inferred without topology constraints and as unrooted trees; rooting was applied only for display or ancestral-state visualization. Alternative placements of A0A951 were evaluated by constrained maximum-likelihood searches compared with the approximately unbiased, KH, SH, and c-ELW tests over 10,000 RELL replicates^31^, and by site-concordance factors^69^; empirical-profile mixture models^70^ were used as sensitivity analyses where noted. Full topology-test results are given in Supplementary Note 2.

##### **Structure-based phylogeny and structural-alphabet comparison**

A structure-based tree was built with FoldMason^71^ over the high-confidence AlphaFold2 and experimental structures, and Foldseek 3Di structural-alphabet strings^10^ were compared directly between the canonical and GHOST sets (Supplementary Note 2).

##### **Historical ancestral-state sensitivity analysis**

Historical sensitivity analysis, not the primary 54-sequence tree: Ancestral states at the avGFP Tyr66-equivalent position were reconstructed by empirical-Bayes marginal reconstruction in IQ-TREE^28^ on the same 49-taxon maximum-likelihood topology and 218-site alignment used for Supplementary Fig. 4, with VT+I+G4 as the primary model. The focal GHOST ancestor, Node N1, was defined as the M-labeled internal node retained from the unpruned 49-taxon topology and shown in Supplementary Fig. 4; broader sensitivity nodes included the nearest canonical cnidarian fluorescent-protein clade. Full ancestral-sequence reconstruction was assessed for robustness across LG^72^, WAG^73^, and LG+C20 profile-mixture models^70^, and under outgroup re-rooting. Sites were called ambiguous when the maximum posterior probability was less than 0.80. The maximum-a-posteriori ancestor used the highest-posterior residue at every retained alignment site, whereas the AltAll uncertainty probe replaced the MAP residue with the second-highest-posterior residue only at sites meeting the posterior-probability ambiguity criterion^74^ (Supplementary Note 2).

##### **Protein structure prediction and quality control**

Structures of family members without experimental models were predicted with AlphaFold2^32^ through ColabFold v1.6.1^33^, using the highest-confidence single-chain model for each protein. Per-residue pLDDT was used for quality control, and every retained model was confirmed to superpose on avGFP with TM-align or US-align. Multi-domain or extension-bearing proteins were handled by retaining the GFP-like barrel region for structural superposition and alignment, while recording terminal extensions or partial coverage as candidate-quality annotations.

##### **Model-ensemble robustness analysis**

For A0A951 and the coral member A0A7U3V473, the five independently predicted, side-chain-relaxed ColabFold models were analyzed as structural ensembles. The R182-E134 salt bridge, the R182 contact to the chromophore-position backbone, the E199-Q197 hydrogen bond, the Tyr83-Tyr168 ring-centroid separation, and the R87-E134 contact were measured in each model. Local pLDDT values and inter-residue predicted aligned error within the core were recorded. fpocket^75^ was also applied across the model ensemble to assess the sensitivity of cavity and druggability assignments to side-chain placement. Distances reported in the Results and Fig. 3 refer to the displayed A0A951 AlphaFold Protein Structure Database model unless otherwise stated. Each model was independently superposed onto the displayed A0A951 AlphaFold model over the GFP-barrel C-alpha atoms. Salt-bridge and hydrogen-bond distances were measured between the minimum functional heavy-atom pairs, whereas Tyr83-Tyr168 and Trp152 aromatic contacts were measured between ring centroids. Robustness was defined by retention of the same contact class across all five models rather than by the shortest distance in a selected model.

##### **ColabFold residue-61 mutational panel**

A0A951HUW2 and the M61V, M61F, M61Y, M61T, and M61H variants were modeled with ColabFold v1.6.1 (commit 0ff4794957abbd5d476a6e1c9b3db853d2b607c9) using the alphafold2_ptm preset and AlphaFold parameter model 1. Multiple-sequence alignments were generated once per construct through the ColabFold MMseqs2 API in mmseqs2_uniref_env mode and reused for both prediction passes. Each construct was evaluated with eight seeds, eight recycles, one ensemble, dropout disabled, and ranking by mean pLDDT. One pass used no templates; the second used 1EMA, 1BFP, and 1ZUX as a fixed custom GFP-family template set, with at most three template hits. No Amber relaxation was performed, yielding 96 unrelaxed structures in total. MSA configuration limits were 512 clustered and 5,120 extra sequences; the server-side database releases were not retained by the ColabFold API.

For each model, minimum heavy-atom distances were calculated between residue 61 and Tyr83, Arg87, Arg182, Gln197, and Glu199, excluding backbone atoms when side-chain atoms were available. Residue-61-to-Tyr83 and residue-61-to-Arg182 distances below 2.4 Å were flagged as short contacts; this threshold was used as a screening rule rather than an energy-based clash definition. Mean pLDDT and within-region PAE were calculated for residues 58-90 and 178-202, with cross-region PAE calculated between the two regions. Barrel closure was summarized by the minimum C-alpha distance and the number of C-alpha contacts below 8 Å between residues 32-43 and 190-201, representing the adjacent first and last barrel strands.

Enclosed empty volume near the pocket was estimated on a 1 Å grid in a 19 × 19 × 19 Å box centered on residues 61, 83, 87, 182, 197, and 199. Element-specific van der Waals radii plus a 1.2 Å probe defined excluded volume; components connected to the box boundary were classified as solvent-accessible, and enclosed components below 3 Å³ were discarded. A transparent compatibility score ranked models within the panel as pLDDT(58-90) + pLDDT(178-202) - 2(cross-region PAE) - short-contact penalty - 0.03(cavity volume) + 0.2(barrel contacts), where the short-contact penalty was 4 × max[0, 2.4 - min(d61-83, d61-182)]. The score was not calibrated to folding free energy, thermodynamic stability, expression, fluorescence, or biochemical activity.

Models were additionally scored against the central-helix criterion of Hartley^83^ and colleagues. For each model the 59 backbone and side-chain atoms of the central-helix segment surrounding the chromophore-equivalent position were superposed by the Kabsch algorithm onto the corresponding atoms of S65T GFP, and the root-mean-square deviation recorded; the published decision threshold separating chromophore-forming from non-forming variants is 1.0487 A. The same procedure was applied to the wild-type models of the other chromophore-position variants in the family and to their tyrosine-substituted designs. Comparisons of this kind are only valid within a single prediction pipeline. Measured across thirteen controls folded by both routes, the AlphaFold-Database-to-ColabFold offset has a median of -0.030 A and a standard deviation of 0.091 A over the eight controls well modelled by both, but individual models can be far worse: one control moved 3.78 A to 0.75 A. Applying the criterion across pipelines changed two of four wild-type verdicts, one from fail to pass and one from pass to fail, so every wild-type and design pair reported here was re-folded by ColabFold and is compared only against same-pipeline values, with 0.091 A taken as the noise floor below which a design shift is not interpreted.

##### **Molecular-dynamics simulation and network-occupancy analysis**

Explicit-solvent molecular-dynamics simulations were run with OpenMM 8.5.2 (OpenCL platform)^36^ using Amber ff14SB^37^ for the protein and rigid TIP3P water^38^. Four systems were simulated: A0A951 with an immature pre-cyclization CIM residue and Glu199 either charged or protonated, and the A0A7U3V473 F67Y coral model with a mature anionic CRO chromophore using the original and a phi-refit chromophore-torsion parameter set. The noncanonical chromophore residues used GAFF2^39^ atom types and AM1-BCC^40^ charges generated from capped fragments with antechamber and parmchk2 in AmberTools 24.8^41^, with explicit junction parameters to the ff14SB flanking residues. Each system was solvated in a rectangular box with 1.0 nm solute padding, neutralized, and brought to 0.15 M NaCl.

Long-range electrostatics were treated with particle-mesh Ewald^76^ using a 1.0 nm real-space cutoff. Bonds to hydrogen were constrained, water molecules were rigid, and the integration timestep was 2 fs. A Langevin middle integrator was used at 300 K with a collision frequency of 1 ps-1, together with a Monte Carlo barostat at 1 bar and 300 K updated every 25 steps. Each replica underwent energy minimization, 0.2 ns NVT equilibration, 1.0 ns NPT equilibration, and 100 ns NPT production; three independent replicas were run per system. Coordinates and state data were saved every 10 ps, yielding 10,000 frames per 100 ns replica.

Trajectories were least-squares aligned on C-alpha atoms and analyzed with MDTraj 1.11.154. Interaction occupancy was defined as the fraction of frames satisfying a distance cutoff of <4.0 Å for cation-anion contacts, <3.5 Å for hydrogen bonds, and <7.0 Å for aromatic ring-centroid contacts. Heavy-atom root-mean-square fluctuations were calculated after C-alpha alignment. Distances are reported with canonical A0A951 or A0A7U3V473 residue numbering; the raw trajectory topologies use numbering shifted by the merged chromophore residue (founder -3; coral -2). Because the founder and coral trajectories represent immature and mature chromophore states, respectively, loss of the coral chromophore-backbone and Gln203/Glu205 cap contacts was interpreted as maturation-dependent rather than as loss of the central R2E2 core. The merged chromophore residue itself was identified directly in each topology rather than by applying the global numbering offset: CIM was topology residue 59 in the founder and CRO was residue 66 in the coral system. The chromophore-backbone contact was measured to the merged residue's tail-carbonyl atom O3 rather than to the generic backbone atom O. Hydration was analyzed on the same trajectories. Pocket hydration was the number of water oxygens within 5 Å of the network residues; per-residue hydration was the fraction of frames with a water oxygen within the 3.5 Å hydrogen-bond cutoff of the residue; and a cation-anion contact was scored as water-bridged in any frame in which a water molecule bridged the two partners. Values were pooled across the three replicas, 30,000 frames per system. The DUF2490 complete-network member A0A8J3CUU4 was simulated under the same protocol in three replicas, and interaction occupancies and per-residue heavy-atom fluctuations were computed as for A0A951 after taking every 20th frame (500 frames per replica).

##### **R2E2 core geometry**

Analyses used the AlphaFold Protein Structure Database model of A0A951 (AF-A0A951HUW2-F1-model_v6) and the AlphaFold/ColabFold family models listed in the master table (mean pLDDT 80 to 97). Interatomic geometry and solvent accessibility were computed with Biopython Bio.PDB 1.82^24^. The interior network comprised Arg87 and Arg182, Glu134 and Glu199, Ser67, Ser146 and Ser166, Gln197, Tyr83 and Tyr168, Trp152, and the Met61 chromophore-position backbone. Minimum functional-atom distances defined salt bridges (<4.0 Å), hydrogen bonds (<3.5 Å), and contacts (<5.0 Å). Aromatic contacts among Tyr83, Tyr168 and Trp152 were assessed from ring-centroid distances, with ≤7 Å treated as a contact, and shared-metal geometry was tested from the Glu134/Glu199 carboxylate-oxygen spacing (3.0 to 7.0 Å bidentate-shell range). Solvent-accessible surface area was calculated by the Shrake-Rupley^34^ algorithm, with <15 Å² classed as buried.

##### **Electrostatic pKa signature across the family**

Titratable-group pKa values were predicted with PROPKA 3.5.1^43^ for A0A951 and each available family model. For every model, the most up-shifted buried arginine (solvent-accessible surface area <15 Å²) was identified as the Arg182-equivalent candidate, its shift was calculated relative to the model arginine pKa of 12.5, and buried acidic counterions were counted.

##### **Continuum electrostatics**

The A0A951 model was protonated and parameterized with PDB2PQR 3.6.1 using the AMBER force field at pH 7.0, and the Poisson-Boltzmann potential was solved with APBS 3.4.1^44^. Electrostatic potential, in units of kT/e, was interpolated at the functional atoms of each network residue and compared with the distribution of C-alpha potentials over the whole protein. Because point potentials at partial-charge-bearing atoms include a self-energy component, they were interpreted qualitatively. This single-founder analysis was run with script 117 using pdb2pqr30 --ff=AMBER --with-ph=7.0 (output a0a.pqr). Its network-wide potentials were described narratively and are not reported in Supplementary Table 10.

In a separate cross-protein analysis, apbs_site_potential.py used PDB2PQR with --ff=PARSE for the 47-structure panel in Supplementary Table 10 (17 GHOSTs, 18 general fluorescent proteins and 12 matched-core carriers). The potential was evaluated at one probe atom per structure, the chromophore-site backbone carbonyl oxygen, with charges on all three chromophore-forming triad residues zeroed. Unlike the AMBER founder calculation, this comparison reports group distributions rather than potentials throughout the network. The computed means were 36.152, 19.442 and 24.771 kT/e, respectively, reported as 36.2, 19.4 and 24.8 kT/e in the table.

##### **Cavity detection**

Enclosed cavities were detected with fpocket 4.2.3 on the displayed A0A951 model and repeated across the five-model ensemble used for robustness analysis. Pockets were ranked by druggability score, and the lining residues of each pocket were intersected with the network core (Arg87, Arg182, Glu134, Glu199, and Tyr168) to test whether a consistently positioned, network-lined cavity was present. Because cavity scores varied with model choice, interpretation emphasized conserved geometry and pocket location rather than any single druggability value. The conda-forge package version was 4.2.3, although the executable startup banner identified itself as fpocket 4.0.

##### **Structural-neighbor and functional-site search**

The A0A951 model was searched against the Protein Data Bank with Foldseek 10.941cd33 easy-search in TM-alignment mode (--alignment-type 1). The 40 top-scoring neighbors were annotated through the RCSB Data API^77^ for entry title and bound non-polymer components, excluding crystallization ions, and classified as GFP-family or other. The Mechanism and Catalytic Site Atlas^78^ was queried for any GFP-fold catalytic entry. An AutoDock Vina^79^ ligand-docking tier was staged for the cofactor/anion hypothesis but was not pursued because no consistently positioned, network-lined open cavity was identified across the model ensemble.

##### **Fold-independent structure-motif searches**

The buried network was defined by A0A951 Arg87, Glu134, Tyr168, Arg182, and Glu199. The RCSB Structure Motif Search service^46^ was queried with the A0A951 AlphaFold model using side-chain atom pairing and RMSD cutoffs from 1.0 to 2.0 Å. One residue-faithful motif allowed Arg/Lys and Glu/Asp exchanges with Tyr held strict; a chemical-class motif allowed cations (Arg, Lys, His), anions (Asp, Glu), and aromatics (Tyr, His, Trp). Searches covered the experimental PDB and the RCSB computed-model collection. Every hit was independently checked for connected side-chain geometry: cation-anion pairs required a charged-nitrogen to carboxylate-oxygen distance below 4.0 Å, and the aromatic group was required within 4.0 Å of a network partner. The PRP21-PRP9 interface hit was cross-checked in PDB entries 4DGW, 5ZWM, and 7DCO and mapped to the human SF3A1 and SF3A3 orthologs by BLOSUM62 alignment.

The AlphaFold database was searched with FoldDisco^47^ through the Foldseek webserver against the AFDB50 cluster-representative index. The chemical motif was submitted as A87:p,A134:n,A168:a,A182:p,A199:n, where p, n, and a denote positively charged, negatively charged, and aromatic residue classes. Full five-position matches were retained and ranked by the FoldDisco rarity score; recovery of known family members at motif RMSD 0.24 to 0.34 Å served as a positive control.

##### **DUF2490 census and convergence analysis**

All 4,234 UniProt members of PF10677 were collected with taxonomic lineage and sequence length. AlphaFold models were available for 2,789 members and were screened de novo by enumerating cationic, anionic, and aromatic side chains and fitting five-residue sets to the A0A951 functional-atom template by Kabsch superposition. A complete R2E2 core required a functional-atom RMSD of at most 1.8 Å, at least two cation-anion salt bridges below 4.0 Å, and aromatic engagement below 4.0 Å. Burial was quantified by relative solvent-accessible surface area using Shrake-Rupley calculations, and model confidence was read from per-residue pLDDT values.

Representative DUF2490 complete-core models were classified by TM-align against avGFP and A0A951, Foldseek easy-search against the PDB100 database, and pydssp secondary-structure assignment. Convergence was assessed by comparing whole-protein TM-scores with superpositions of the five network residues alone, reported as side-chain functional-atom and C-alpha RMSDs. The all-versus-all TM-align/Seq_ID matrix used a separate TM-align build from the US-align v20260527 executable used to establish the avGFP residue correspondences.

##### **Predicted subcellular localization**

Signal peptides were predicted with SignalP 6.0^48^ in fast mode with organism class 'other', run remotely through the DTU SignalP 6 application on BioLib. The test set comprised seven retrievable GHOST sequences and a random sample of 60 DUF2490 sequences. Predictions were summarized by class (OTHER or Sec/SPI) and cleavage-site position.

##### **Structural-figure rendering**

Fig. 1 and Supplementary Fig. 3a,b were rendered with PyMOL 3.1.0 (open-source, conda-forge) from Python scripts and ray-traced on a white background. Residue labels and leader lines were added afterwards in Matplotlib: the screen position of each labeled atom was obtained by ray-tracing that atom alone as a pseudoatom in the same view, and labels were placed in the margins so that none overlaps the structure. For Supplementary Fig. 3, A0A8J3CUU4 was superposed on A0A951 with pair_fit over the five role-matched functional-atom pairs (Arg CZ, Glu CD, Tyr OH), giving a functional-atom RMSD of 1.20 Å; this display alignment was separate from the TM-align/US-align calculations used for quantitative structural relatedness.

##### **Across-family conservation of the R2E2 core**

Conservation of each network position was evaluated across the 22 full-length family members at the sequence level. Positions in the validated 21-residue diagnostic fingerprint, including Ser67, Arg87, Glu134, Trp152, Tyr168, Arg182, Gln197 and Glu199, were taken from the fingerprint column of the master table; Tyr83, Ser146 and Ser166 were read from the 25-sequence, 327-column family alignment mapped through A0A951. Chemical character was scored with conservative substitutions allowed for cations (Arg/Lys), anions (Glu/Asp), amides (Gln/Asn), and hydroxyl residues (Ser/Thr); the putative redox position required tyrosine.

##### **ConSurf and maximum-likelihood site-specific evolutionary-rate analysis**

GHOST sequence conservation was calculated with the ConSurf server62 using an external, query-anchored multiple sequence alignment rather than server-selected homologs. The completed ConSurf run mapped 218 A0A951 positions from 28 aligned sequences and assigned conservation grades with the ConSurf^45^/Rate4Site^80^ implementation. A separate local enrichment analysis used Shannon-entropy conservation scores from the bacterial/GHOST alignment and mapped them onto the A0A951 AlphaFold model. Solvent exposure was calculated with Shrake-Rupley solvent-accessible surface area; residues with relative SASA less than 0.10 were classified as buried/internal, and residues with relative SASA greater than 0.25 as surface-exposed. The chromophore/network neighborhood was defined by the structurally mapped active-site regions and residues within 8 angstroms of the G62, P64, S67, Y83, R87, E134, W152, P177, R182, Q197 and E199 landmarks. Conservation enrichment was assessed with 20,000 random permutations and a Mann-Whitney U test. The network-neighborhood and surface groups contained 55 and 72 residues, respectively; the observed mean conservation difference was 0.186, Mann-Whitney U = 3025.5, one-sided P = 1.83 x 10-7, and the rank-biserial effect size was 0.528.

For the analysis shown in Fig. 5, site-specific evolutionary rates were estimated as empirical-Bayes posterior means under LG with a 20-category discrete-Gamma rate distribution, with the topology fixed to the maximum-likelihood tree of the 25-sequence family alignment. That alignment is an A0A951-anchored MAFFT L-INS-i^26^ alignment of 325 columns over the corrected roster, in which the redundant gene model A0A146F2F0 was withdrawn and A0ABR1SEL9 added; the tree was inferred in IQ-TREE 3 under LG+I+G4 selected by BIC. The sequence set comprised all 22 complete barrels and the three partial fragments RKT55308, HEY8628922, and MEY2932110. Alignment gaps and nonstandard characters were treated as missing data, so partial sequences contributed missing data outside their aligned spans. No gap-occupancy threshold was applied during likelihood calculation. For Fig. 5 and the core-vs-barrel tests, rates were retained for the 218 alignment columns corresponding to A0A951 residues; insertion-only columns without an A0A951 coordinate were excluded. The earlier record of this analysis as an LG+G4 run on a 327-column FAMSA alignment of the superseded roster has been replaced by the regenerated inference described here.

Posterior-mean rates were reported on the standard scale on which the discrete-Gamma categories have a mean of 1; no additional across-site normalization was applied. Twenty rate categories were used rather than four, because the four-category model places every slow site at the same category floor and so cannot resolve the core positions either from one another or from the chromophore site. Percentiles were calculated by ranking the 218 A0A951-mapped columns from lowest to highest rate using ordinal ranks, with ties broken by residue order. Lower percentiles therefore indicate slower evolution and stronger constraint. The primary functional test, confirmatory rather than prespecified, compared the five-residue charged/redox core (R87, R182, E134, E199, and Y168; n = 5) with the remaining 213 mapped barrel positions using a one-sided Mann-Whitney U test (alternative hypothesis: core rates are lower; U = 12, p = 9.58 × 10−5), with a two-sided value reported as a sensitivity analysis (p = 1.92 × 10−4). Adding the Q197 cap as a sixth core position gave a one-sided p = 2.26 × 10−5. A separate diagnostic-position test used P64, S67, E134, W152, R182, and Q197 (one-sided p = 4.58 × 10−5) and was interpreted as a family-identity rather than a mechanistic comparison. The separate ConSurf^45^ analysis used a 28-sequence broad-context alignment, whereas the maximum-likelihood rate analysis and sequence logo used the final 25-sequence family alignment. The 28-versus-25 difference therefore reflects distinct input alignments and not exclusion of the three partial fragments, which were retained in the 25-sequence analysis. To separate constraint from burial, relative solvent accessibility was computed for every residue of the A0A951 AlphaFold model with the Shrake-Rupley algorithm in Biopython, normalized to the theoretical maximum accessibility of each residue type (Tien et al. 2013) ⁹². The five core rates were compared with those of all other positions at or below accessibility thresholds of 0.01, 0.05, 0.10 and 0.20 by one-sided Mann-Whitney U test, and by a permutation test drawing 100,000 random sets of five positions from each buried pool. The core is significant at every threshold (Supplementary Data).

##### **Chromophore and buried-network fingerprint; two-group sequence logo**

Because sequence identity between the GHOST family and canonical fluorescent proteins is low (23 to 35% to avGFP over aligned residue pairs; Supplementary Table 11), a full-length between-group multiple alignment is unreliable; therefore, the diagnostic positions were matched by structural equivalence (TM-align^21^ and US-align^22^) rather than by sequence alignment. The two-group sequence logo (Fig. 2) was drawn with Logomaker^81^ over nine structurally defined positions: the conserved Gly31/Gly33/Gly35 scaffold positions (A0A951 Gly27/Gly29/Gly31), the Tyr66-equivalent site (Met61), the triad Gly67 (Gly62), the catalytic Arg96 and Glu222 (Arg87 and Glu199), and the pocket positions Leu201 and Leu220 (Arg182 and Gln197). Family residue frequencies were taken from the 25-sequence query-anchored A0A951 GHOST alignment, and canonical fluorescent-protein frequencies from a curated FPbase-derived wild-type-like Tyr66 GFP-family fluorescent-protein panel of 349 sequences, each pairwise-aligned to avGFP by a BLOSUM62^23^ global alignment (gap-open penalty -10; gap-extension penalty -0.5) in Biopython^24^ to read the residue at every avGFP position. These global alignments were used only to project the predefined, structurally equivalent avGFP positions onto the broad fluorescent-protein panel. The separate 14-protein structural panel was used for local pocket and structural comparisons and did not contribute to the sequence-logo frequencies.

##### **Genomic context and mobile-element analysis**

The source contig (JAFAZW010000077.1) and the genomic neighborhoods of eleven further homologs were retrieved from their source assemblies. Neighborhoods were analyzed over approximately eight genes on either side of each GHOST locus when contig length permitted. Open reading frames were called concordantly with the NCBI Prokaryotic Genome Annotation Pipeline66 and Prodigal^50^/Pyrodigal^51^; physical contiguity of the source locus was confirmed by mapping the source-metagenome reads with BWA-MEM^52^ and SAMtools^53^. Mobile genetic elements and viral segments were classified with geNomad^54^, CheckV^55^ and VirSorter2^56^. Neighborhoods were manually reviewed for att-site-like repeats, direct repeats, tRNA insertion sites, integrases, recombinases, transposases, relaxases, oriT-like motifs, conjugation genes, toxin-antitoxin modules, restriction-modification genes, Abi/PDC-like defense genes, and local GC or codon-usage shifts. Per-gene GC content and a sliding-window GC scan were computed across each locus to detect the compositional GC-island signature of horizontal transfer (Supplementary Note 1).

##### **Defence-system screen**

Anti-phage defence systems in the twelve neighborhoods were called with two independent annotators, DefenseFinder^57^ and PADLOC^58^. Defence-adjacency of the barrel gene was tested against length-matched genes drawn from the same assemblies by permutation, with the null distribution restricted to genes from the same local genomic contexts rather than whole-genome backgrounds. An explicit anti-defence or counter-restriction screen was then applied by searching neighboring annotations for ArdA-like proteins, restriction-modification systems, mobile-element recombinases, toxin-antitoxin modules and other markers of mobile-element self-protection (Supplementary Note 1). Control proteins were matched within 0.70-1.30 times the focal protein length when possible, genes already annotated as defense or mobile-element markers were excluded from the control pool, and feature adjacency was scored within +/-10 genes. Source-habitat comparisons were coded separately because same-assembly controls necessarily share the habitat metadata of the focal locus.

##### **Ecological analysis**

For each of the 22 full-length members, source biome, redox/irradiance context, biological origin, and an extreme-niche flag (polar, thermal, desiccation/ultraviolet, or high-irradiance reef) were derived from the collection country (NCBI BioSample geo_loc_name) and ecosystem recorded in the per-member master table (01_query_and_models/A0A951HUW2_master_table.md in the archived data set). Records were grouped as surface soil, sub-oxic groundwater, thermal spring, other marine, coral reef, terrestrial fungi, or lichen. Biological origin was classified as free-living environmental microbe, host-organism-encoded, or host/symbiont-associated. Habitat comparisons were treated as descriptive because the structure/profile discovery pipeline does not provide a matched database-wide ecological denominator and the sequences are phylogenetically non-independent.

##### **Statistics and reproducibility**

Software versions are given with each analysis above; statistical conventions are stated in the main Methods. Function analyses used Biopython 1.82, PROPKA 3.5.1, SciPy 1.13.177 and NumPy 2.0.278. Molecular-dynamics production ran on two NVIDIA Quadro RTX 5000 GPUs under Ubuntu 26.04 LTS (NVIDIA driver 580.159.03). Exact command lines and custom analysis code are in the scripts/ folder of the archived data set (Data availability). The date and client version of the historical remote BLAST search were not recoverable and should be recorded in any confirmatory rerun. The GOLD^82^ binomial calculation is retained only as an illustrative heuristic and is not interpreted as a formal ecological enrichment test. Quantum-chemical characterization of the buried network is reported in the companion study.

#### **Supplementary Note 1 | Provenance, genomic context, and defense analyses**

##### **S1.1 Founding-contig provenance**

A0A951 is encoded on contig JAFAZW010000077.1 from a Brazilian rhizosphere-soil metagenome. The contig is 89% coding at 1.04 genes per kb, carries an upstream bacterial Shine-Dalgarno motif, and is flanked by bacterial and phage-associated genes without eukaryotic markers. NCBI PGAP and Prodigal return the same open reading frame and start site. Mapping SRA run SRR8113284 shows continuous coverage, paired fragments spanning both gene boundaries, and no boundary-associated coverage cliff or soft-clip pileup. geNomad assigns probabilities of 0.94 chromosome, 0.04 plasmid, and 0.02 virus and calls no provirus; VirSorter2 returns no viral sequence, and CheckV reports no provirus. These independent tests support a genuine bacterial chromosomal locus rather than a binning artifact or synthetic construct.

##### **S1.2 Source cassette and GC island**

A0A951 is the only structurally characterized member of a five-gene, same-strand cassette. The cassette is approximately 57% GC against a 71% contig median, a roughly 13-percentage-point drop that falls at the sixth percentile and returns to chromosome-like composition on both flanks. It lies adjacent to a BrnT/BrnA toxin-antitoxin module and a phage-type DNA-packaging gene. The compositional discontinuity is consistent with horizontal acquisition, although geNomad, VirSorter2, and CheckV identify no intact integrated element.

##### **S1.3 Cross-member neighborhoods**

The flanking genes of eleven additional homologs differ almost completely among taxa, arguing against a conserved operon and favoring a mobile accessory locus. In every examined high-GC host, the family gene lies in a local 4-10 percentage-point GC island, whereas in low-GC hosts it generally matches host composition. In the best-assembled genomes, the locus repeatedly occurs within approximately eight genes of mobile-element or anti-phage machinery. The genomic context of the founder locus is shown in Supplementary Fig. 2.

##### **S1.4 Defense-system and anti-defense tests**

DefenseFinder and PADLOC never place a GHOST gene within a called defense system. PADLOC nevertheless detects bona fide neighboring systems in two unrelated phyla: a type I restriction-modification system with PDC-S13 in one deltaproteobacterial neighborhood and a type I restriction-modification system with AbiE in a Bacteroidota neighborhood, each approximately eight genes from the barrel gene. Against length-matched genes from the same assemblies, the family gene is not significantly more defense-adjacent than neighboring genes (permutation p approximately 0.1). One *Maribacter* locus lies near an ArdA anti-restriction gene and recombinases, but the family proteins lack the strongly acidic, DNA-mimicking surface expected for classical anti-restriction proteins. The supported interpretation is therefore a horizontally mobile accessory locus embedded in mobile-element-rich neighborhoods, not a defense or anti-defense component.

A matched-window analysis compared 12 GHOST loci with retrievable genomic windows against 103 length-matched, non-marker control proteins from the same windows. The family loci were not enriched for defense adjacency (2/12 focal loci versus 28/103 controls; one-sided p = 0.875) or for combined defense/mobile adjacency (4/12 versus 35/103; p = 0.633). Mobile-element proximity alone was suggestive but not significant (2/12 versus 7/103; one-sided p = 0.237; paired permutation p = 0.0649). These results support the cautious interpretation of occasional mobile-context proximity without evidence that the barrel itself is a defense-system component.

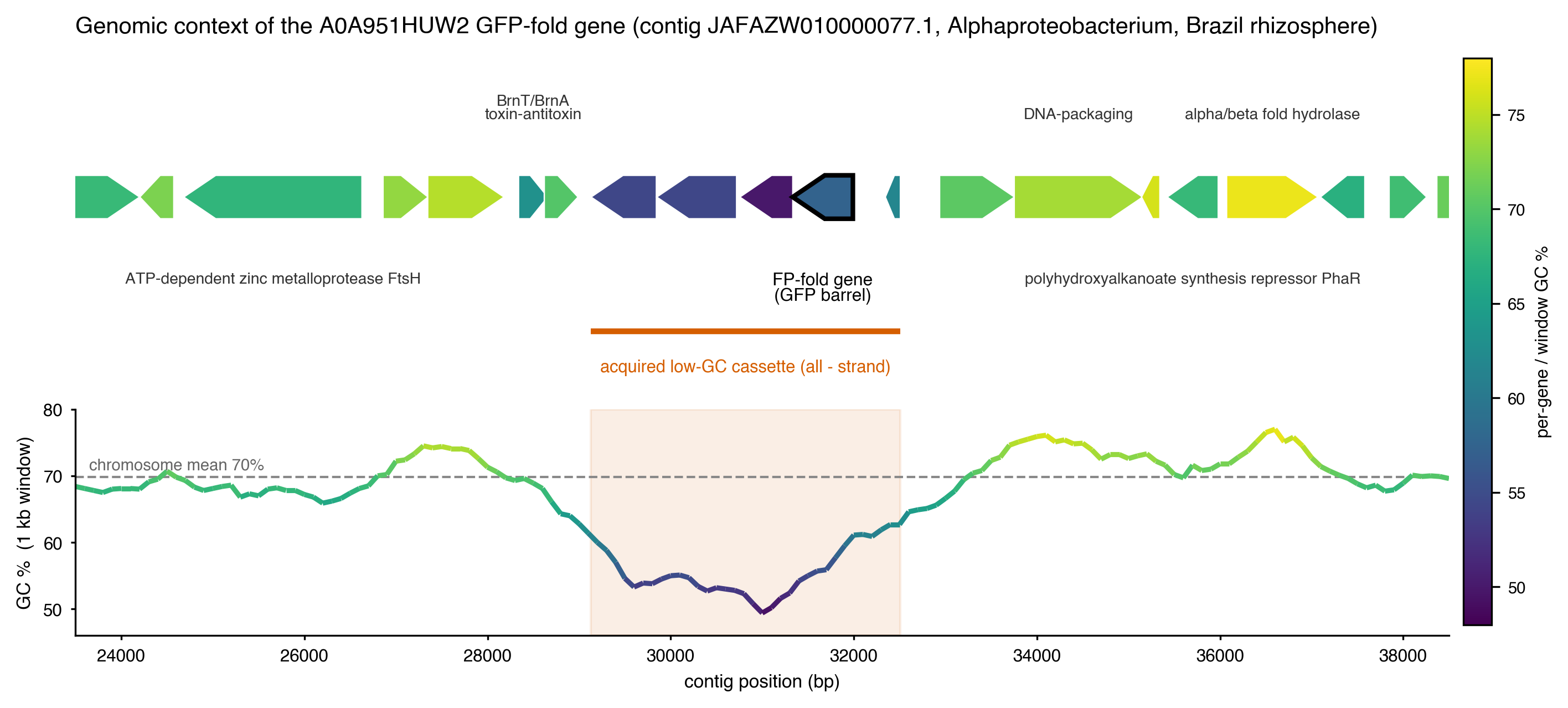

Supplementary Fig. 2 | Genomic context of the founder GHOST gene. Gene map of contig JAFAZW010000077.1 (Alphaproteobacteria metagenome-assembled genome, Brazilian rhizosphere soil) around A0A951HUW2 (outlined), with genes coloured by GC content and a 1-kb sliding-window GC profile below. The GHOST gene lies in a five-gene, same-strand cassette (orange bar) whose GC content falls to about 57%, against about 70% for the surrounding chromosome (dashed line). The cassette is flanked by a BrnT/BrnA toxin-antitoxin module and a phage-type DNA-packaging gene. Comparisons across eleven further GHOST loci are given in Supplementary Note 1 (S1.3).

#### **Supplementary Note 2 | Family boundary, phylogenetic placement, and ancestral robustness**

##### **S2.1 Profile-HMM calibration and database saturation**

A profile hidden Markov model built from ten representative members cleanly separates the GHOST family from canonical fluorescent proteins: family members score 233-309 bits, the highest-scoring canonical fluorescent protein scores 82 bits, and none of 990 FPbase proteins exceeds the 157-bit inclusion threshold. Searches of UniProtKB and MGnify add only one metagenomic bacterial sequence and a small set of fungal barrels before saturating, with no long tail of additional homologs. The family is therefore a discrete and genuinely rare GFP-fold lineage rather than a diffuse, undersampled edge of the canonical fluorescent-protein family.

The 157-bit threshold was selected as a high-confidence expansion threshold, not as an absolute family boundary. In leave-one-out tests of the ten-member bacterial seed, the held-out seed scores ranged from 120.6 to 175.6 bits, while the strongest FPbase/Swiss-Prot control score was 155.9 bits. Placing the threshold just above that control reduced false inclusion of canonical fluorescent proteins, but also meant that divergent seed-like sequences below 157 bits were evaluated only as sensitivity candidates. The complete-membership claim therefore combines profile score with full-barrel coverage, the GHOST fingerprint, structure, and provenance rather than relying on the threshold alone.

##### **S2.2 EGFP-free topology tests**

Topology tests were performed after excluding three reference assemblies containing EGFP-like sequences attributable to synthetic-construct contamination. Relative to the unconstrained bacterial placement, forcing A0A951 inside the canonical fluorescent-protein clade gives Δlog L = 1424.8 and p(AU) = 3.54 × 10−40, reported in the manuscript as 3.5 × 10−40. Forcing A0A951 to be sister to the *Acropora* sequences gives Δlog L = 75.9 and p(AU) = 1.64 × 10−6. The unconstrained bacterial placement is the only tested topology not rejected. KH, SH, and c-ELW tests over 10,000 RELL replicates agree. A superseded pre-cleanup analysis that retained the EGFP-contaminated assemblies yielded a more extreme p(AU) value and is not used in the manuscript.

Site-concordance factors are modest (mean approximately 35%), as expected for a deep, low-identity family in which resampling support can exceed site-level concordance. The placement is therefore based principally on the topology tests rather than on bootstrap values alone. Four sequences, A0A951HUW2, MBV9931914, RKT55308, and XWJ43021, show significant amino-acid composition heterogeneity; placements involving the focal and *Saccharothrix*-like branches should be interpreted with this sensitivity in mind. The complete 49-taxon branch-length phylogram is provided as Supplementary Fig. 4.

##### **S2.3 Structure-based and 3Di comparisons**

FoldMason has limited resolving power because the global GFP barrel is strongly conserved and canonical and GHOST structures can interleave. Direct Foldseek 3Di comparisons nevertheless show lower mean distances within canonical fluorescent proteins (0.34) and within the bacterial/A0A951-like set (0.43) than between the groups (0.54). Around the chromophore helix and buried network, the *Acropora* phenylalanine protein is closer to the bacterial set than to canonical fluorescent proteins (3Di distance 0.39 versus 0.46), with 7 of 10 local positions matching the bacterial state. These structural-alphabet results support, but do not independently determine, the sequence-based placement.

As a further robustness check, seven nidogen G2 beta-barrel domains, including PDB 1H4U, were added to the 50-taxon GFP-like sensitivity set. The 57-sequence IQ-TREE 3.1.3 analysis under LG+G4 placed the nidogens in a single, very distant group, with the nearest patristic distance to a GFP-like sequence approximately four substitutions per site. The nidogen attachment was weakly supported and composition-flagged, as expected for a remote barrel analogue. Their inclusion did not move A0A951HUW2, whose nearest neighbors remained bacterial and *Acropora* proteins. Nidogen was therefore treated as a distant structural analogue rather than a close homolog and was not used to root the main-text tree.

##### **S2.4 Ancestral-sequence reconstruction robustness**

Historical sensitivity analysis, not the primary 54-sequence tree: The methionine call at Node N1 and the ancestral R2E2 core are recovered under VT+I+G4, LG, WAG, and LG+C20 sensitivity analyses, under outgroup re-rooting, and in both maximum-a-posteriori and AltAll ancestors. The maximum-a-posteriori and AltAll sequences differ at 91 of 227 alignment positions, all at fast-evolving surface sites, but at none of the chromophore, catalytic, or R2E2 core positions. Thus, the ancestral non-tyrosine chromophore site and buried network are not artifacts of a single substitution model or ancestral-sequence choice.

##### **S2.5 *Acropora*-specific sequence affinity**

Tyrosine-containing *Acropora* GFP-like proteins have a median identity of 24.8% to A0A951, only marginally above the 23.9% median for the wider fluorescent-protein set. In contrast, the two phenylalanine-containing eqFP611-like *Acropora* proteins share 39-42% identity with A0A951, exceeding every one of approximately 1,200 fluorescent proteins examined. Their bacterial affinity is therefore specific to the rare GHOST proteins rather than a general property of *Acropora* fluorescent proteins.

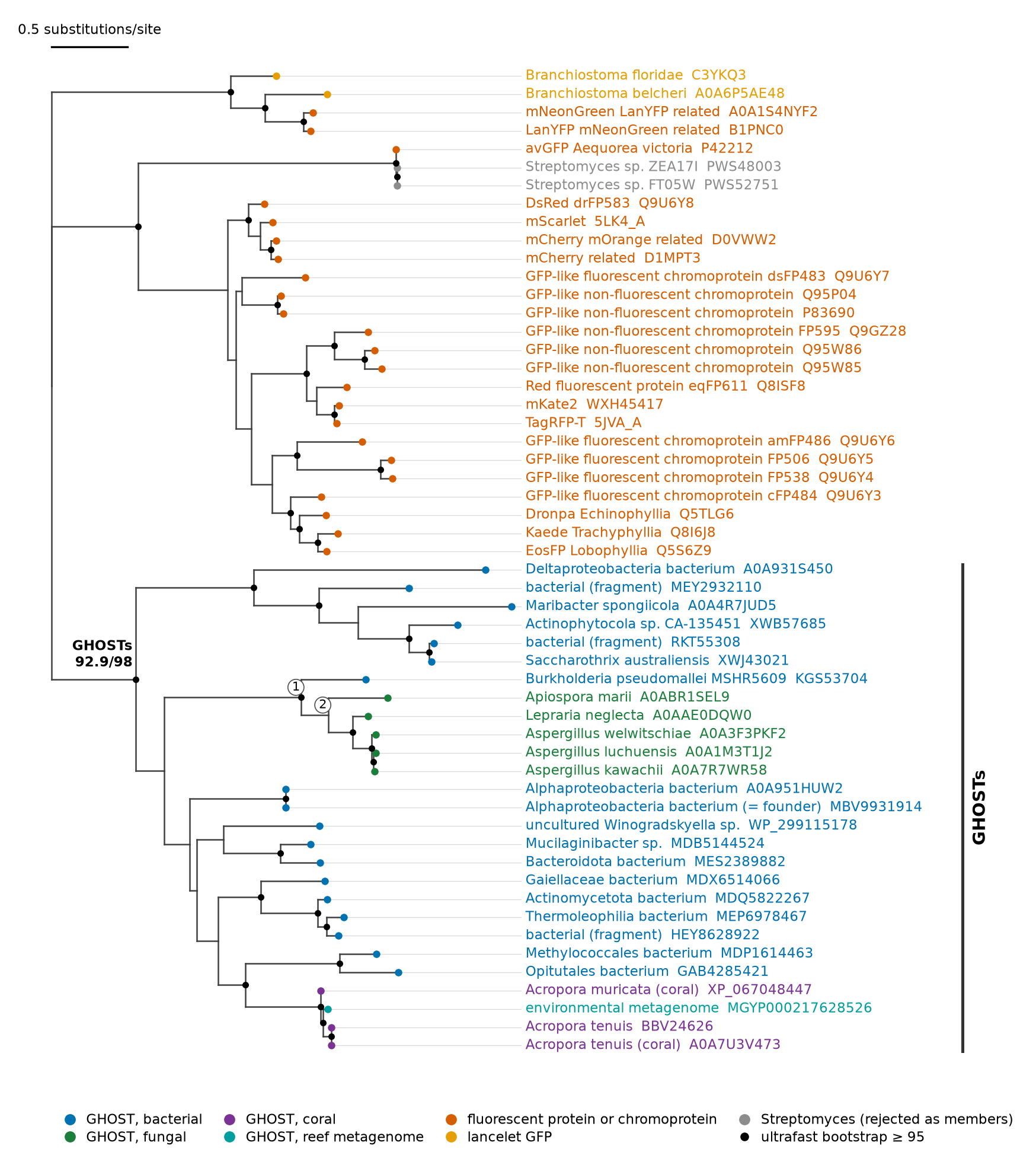

Supplementary Fig. 1 | Maximum-likelihood phylogeny of the GHOST family and GFP-family reference proteins. Tree of 54 sequences (MAFFT L-INS-i alignment trimmed with trimAl^27^ at a gap threshold of 0.2, 228 columns) inferred in IQ-TREE 3 under LG+G4 selected by BIC. The tree is unrooted and is drawn rooted on the fluorescent proteins, chromoproteins and lancelet GFPs only for display. Support values are SH-aLRT / ultrafast bootstrap (1,000 replicates each). They belong to bipartitions of the unrooted tree and were mapped onto the display by split rather than carried with node labels. Black dots mark ultrafast bootstrap of 95 or more. All 25 GHOST records form one clade (92.9/98), together with MBV9931914 (identical to the founder) and BBV24626 (a second *Acropora tenuis* entry). Node 1: the five fungal GHOSTs with KGS53704 (100/100). Node 2: the five fungal GHOSTs (80.7/85; 94.4/97 without the long-branched A0ABR1SEL9). The two Streptomyces sequences carried in earlier versions of the roster group with avGFP and are rejected as GHOST members by the approximately unbiased test. Scale bar, 0.5 substitutions per site.

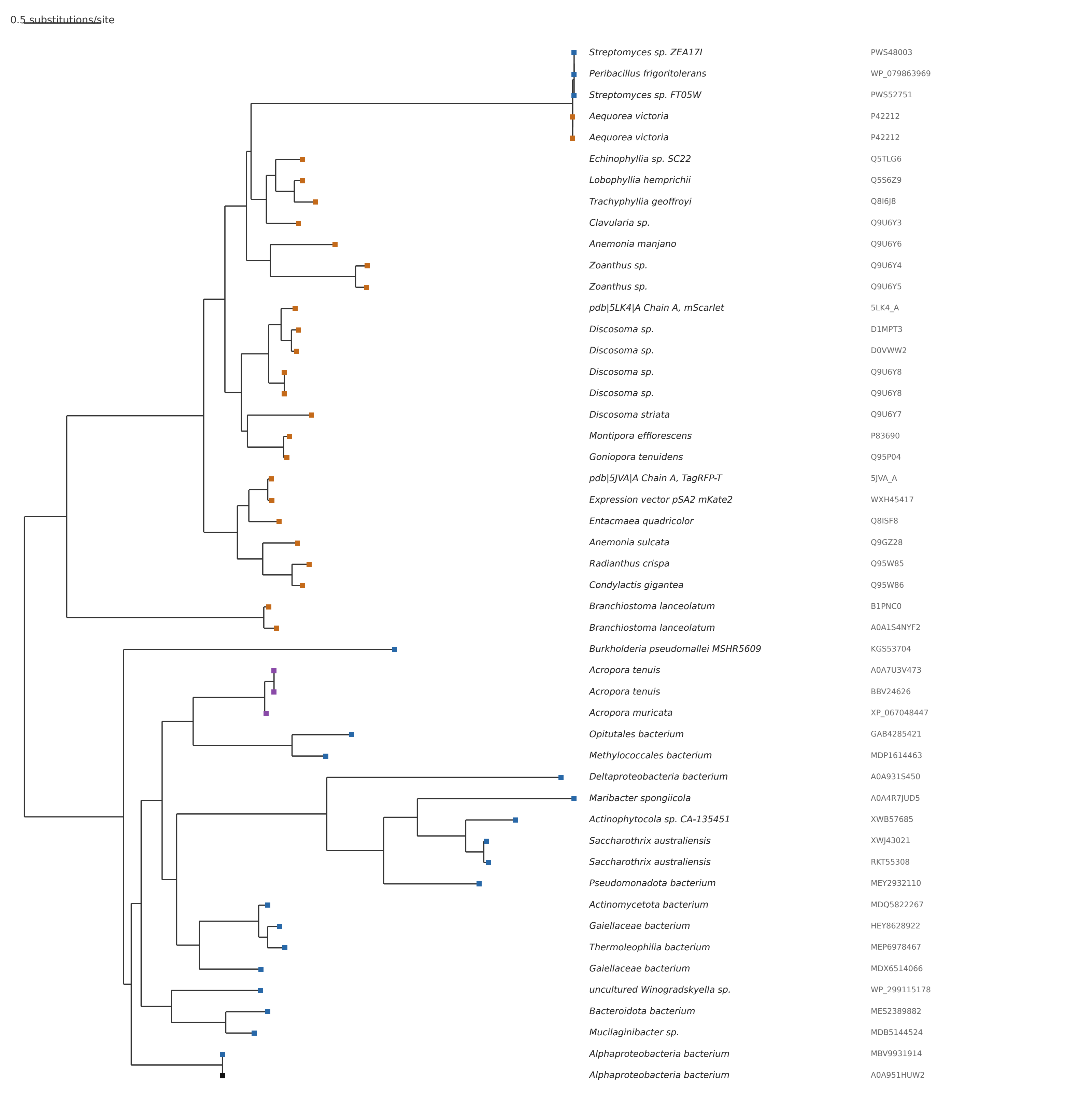

Supplementary Fig. 4 | Historical 49-taxon cross-domain phylogeny. Complete branch-length maximum-likelihood phylogram from the earlier 49-sequence, 227-site analysis under LG+I+G4 (IQ-TREE 3.1.3). It is retained because the historical ancestral-state sensitivity analysis, including Node N1, was run on this topology. It predates the roster corrections: it contains no fungal GHOSTs, includes the two Streptomyces sequences now rejected as members, includes Peribacillus frigoritolerans WP_079863969, which sits with avGFP at near-zero distance, and lists avGFP twice. The primary phylogeny is Supplementary Fig. 1.

#### **Supplementary Note 3 | Conserved glycines of the GFP scaffold**

the GHOST family preserves the Gly31, Gly33, and Gly35 scaffold signature of GFP-like proteins. Gly35 is invariant across the complete family, and Gly31 is retained in nearly all members. Gly33 alone shows a localized relaxation, remaining glycine in about 70% of family members compared with roughly 90-95% in the *Acropora* GFP-like proteins and canonical fluorescent proteins; several bacterial members carry serine at this position. The family therefore retains the structural glycine constraints characteristic of the GFP barrel while tolerating a single localized substitution pattern at Gly33.

#### **Supplementary Note 4 | Structural robustness of the buried R2E2 core**

To test whether the R2E2 core geometry depended on one AlphaFold side-chain placement, we analyzed five independently predicted, side-chain-relaxed ColabFold models spanning A0A951 and the coral GHOST member A0A7U3V473. In all models, the R182-E134 salt bridge, the R182 contact to the chromophore-position backbone, the E199-Q197 hydrogen bond, and the Tyr83-Tyr168 aromatic pair were retained. Principal contact distances varied by less than approximately 0.2 A. The R87-E134 distance was more model-dependent and is best interpreted as a longer-range electrostatic contribution rather than a strict salt bridge. Network residues had local pLDDT values of approximately 95-99 and low inter-residue predicted aligned error within the core. The conclusion therefore rests on retention of contact classes across the ensemble, not on optimization of one selected model.

#### **Supplementary Note 5 | Explicit-solvent molecular-dynamics stability of the buried R2E2 core**

To test the R2E2 core beyond static structure predictions, we analyzed explicit-solvent molecular-dynamics trajectories for the founder A0A951 and the coral member A0A7U3V473. Four systems were examined, each in three independent 100-ns replicas: A0A951 with an immature pre-cyclization CIM residue and Glu199 either charged or protonated, and A0A7U3V473 F67Y with a mature anionic CRO chromophore using the original and a phi-refit chromophore-torsion parameter set.

In A0A951, the R182-E134 and R87-E134 cation-anion contacts were present in 100% of frames in every replica and in both Glu199 protonation states, with mean distances of 2.71 and 2.78-2.79 Å, respectively. The R182-M61 backbone contact and R87-S166 hydrogen bond were also occupied in 100% of frames, and the E199-Q197 cap in 97-100%. The R87-Tyr168 hub hydrogen bond (78-80%) and Tyr83-Tyr168 aromatic contact (59-67%) were more dynamic. Heavy-atom RMSF values of 0.44-0.77 Å for the network residues further indicate a rigid, preorganized founder core. Protonating Glu199 did not materially change these occupancies or distances. In the same trajectories the bridging-water fraction was 0.00 for both cation-anion contacts, per-residue hydration was 0.00-0.02 for Arg87, Arg182 and Glu134 and 0.00 for the Met61 carbonyl, and Glu199 was hydrated in 0.98 and 0.92 of frames with Glu199 charged and protonated respectively; the network shell held 2.6 ± 1.2 and 3.1 ± 1.8 water oxygens within 5 Å in those two states.

In the mature F67Y coral models, the homologous R188-E140 salt bridge remained occupied in 100% of frames, and the R93-S172 contact was retained in 88-99%. The R93-E140 catalytic-arginine arm was more variable (41-47%), while the Tyr89-Tyr174 aromatic contact was present in 16-31%. The R188-chromophore-backbone and E205-Q203 cap contacts were not maintained in the mature state, consistent with chromophore-associated reorganization of the backbone and cap region. Thus, the central di-arginine/bridging-glutamate architecture is the dynamically stable heart of the R2E2 core, whereas peripheral contacts and the mature chromophore-facing region remain more mobile. These results establish that the buried charge cluster is a preorganized structural feature rather than an artifact of one predicted side-chain arrangement, while not identifying a substrate or reaction. The founder simulations represent an immature pre-cyclization state, whereas the coral simulations represent a mature chromophore state. The network residues themselves are standard amino acids, but the chromophore-facing backbone and cap contacts should therefore be interpreted as maturation-dependent.

#### **Supplementary Note 6 | Cavity and functional-site negative controls**

Cavity calls were sensitive to side-chain placement. fpocket best-pocket druggability scores varied from 0.16 to 0.86 across the model ensemble, and no consistently positioned, network-lined open cavity was detected. Functional interpretation therefore emphasizes conserved burial and geometry rather than one predicted cavity. The A0A951 AlphaFold model was also searched against the Protein Data Bank with Foldseek easy-search in TM-alignment mode. The 40 closest structural neighbors were annotated from RCSB entry titles and bound non-polymer components, with ions and common crystallization additives excluded from functional interpretation. All 40 were GFP-family proteins. A query of the Mechanism and Catalytic Site Atlas recovered no GFP-fold catalytic mechanism. These negative controls support a previously unrecognized buried electrostatic/redox architecture while leaving its specific chemical activity unresolved. These nearest-neighbor and M-CSA negative controls do not imply that the residue constellation is unique to the GFP fold; the fold-independent search and convergent DUF2490 recurrence are described in Supplementary Note 7.

#### **Supplementary Note 7 | Fold-independent search for the buried R2E2 core and its convergent recurrence in DUF2490**

##### **S7.1 Structure-motif search of the PDB and computed models**

Seeded on A0A951 Arg87, Glu134, Tyr168, Arg182, and Glu199, the RCSB Structure Motif Search^46^ was run with conservative charge exchanges and, separately, as a chemical-class motif. Three PDB entries matched at RMSD ≤2.0 Å (4NSL, 1.69 Å; 6G4W, 1.78 Å; 6G4S, 1.95 Å), and none matched below 1.5 Å. All three were C-alpha-backbone coincidences: the equivalently typed side chains were 9.6 to 27.3 Å apart and formed no network. Searching the 1,062,058-model RCSB computed set returned one additional chemical-class hit, A2AXG5 at 1.99 Å, which was likewise a backbone coincidence. A direct connectivity check therefore found no self-contained non-GFP R2E2 core in these collections.

##### **S7.2 The PRP21-PRP9 spliceosome interface**

The only non-GFP experimental site reproducing a genuine salt-bridged cluster was the yeast SF3a interface between PRP21 and PRP9. PRP9 Glu5 bridges PRP21 Lys97 (2.9 Å) and Arg101 (3.3 Å) and hydrogen-bonds Tyr142 (2.5 Å). The contacts are present in the crystal structure 4DGW and cryo-EM states 5ZWM and 7DCO. The site differs categorically from the GHOST R2E2 core: it is intermolecular, only partially buried, and matches A0A951 at functional-atom RMSD 2.48 Å despite a C-alpha RMSD of 0.86 Å. It is also not conserved in the human orthologs: SF3A1 retains the Lys and Tyr equivalents but the second cation is Gln, and the PRP9 N-terminal bridging glutamate is replaced by Leu in SF3A3.

##### **S7.3 FoldDisco search of AFDB50**

FoldDisco^47^ was run over the AFDB50 cluster-representative index, approximately 53 million models, with the network encoded as A87:p,A134:n,A168:a,A182:p,A199:n. Known GHOST proteins were the top complete matches at motif RMSD 0.24 to 0.34 Å. Among 107 complete five-position matches, one additional coral GFP-fold candidate from Pocillopora damicornis (A0A3M6US03) was recovered. Because this sequence has not yet undergone the full provenance and family-boundary audit applied to the curated set, it was not added to the 22-member census. Most remaining novel matches belonged to DUF2490 (PF10677).

##### **S7.4 Full DUF2490 census**

DUF2490 comprises 4,234 UniProt members, 84% Bacteroidota and 14% Pseudomonadota, with a median length of 237 residues and no experimental structure in the PDB. Of 2,789 members with an AlphaFold model, 698 (25%) contained the two-cation/two-anion/tyrosine constellation. Twenty-eight had no salt bridge among the matched residues, 421 had one, 222 had two, and 27 had three. A total of 238 members met the complete-core criterion of at least two salt bridges with the aromatic engaged below 4.0 Å. In representative complete-core members, the network was buried (mean relative solvent accessibility 0.04 to 0.05) and modeled at high confidence (local pLDDT 96 to 99; whole-model mean approximately 87). The complete list is archived as 11_motif_search_convergence/census/DUF2490_complete_core_members.tsv.

##### **S7.5 DUF2490 is an outer-membrane beta-barrel**

Foldseek classification of a complete-core DUF2490 model against PDB100 returned outer-membrane beta-barrels, including NanC porin, LpxR, and copper- and polysaccharide-transport barrels, but no GFP-fold structure. DUF2490 lacks the central chromophore-bearing helix, is larger than the GFP barrel, and superposes on avGFP and A0A951 at TM-score 0.34 to 0.40 over approximately half its length, with approximately 5% sequence identity. Superposition of the five network residues alone gave a side-chain functional-atom RMSD of 1.2 to 1.35 Å while the corresponding C-alpha atoms differed by approximately 3.0 Å. An all-versus-all TM-align comparison across canonical fluorescent proteins, GHOST members, and DUF2490 confirmed that the family is FP-related rather than DUF-derived: A0A951 superimposed on canonical fluorescent proteins at TM-score 0.862 (Seq_ID 22%) and on family members at 0.870 (Seq_ID 31%), but on DUF2490 at only 0.365 (Seq_ID 7%). Average-linkage clustering on structural distance placed all canonical fluorescent proteins and GHOST members in one cluster, with DUF2490 as a separate cluster (11_motif_search_convergence/relatedness_TM_seqid.md). The shared network is therefore a convergent active-site-like arrangement rather than evidence of common ancestry with DUF2490, although very deep undetectable homology cannot be formally excluded. The network-focused structural overlay is shown in Supplementary Fig. 3.

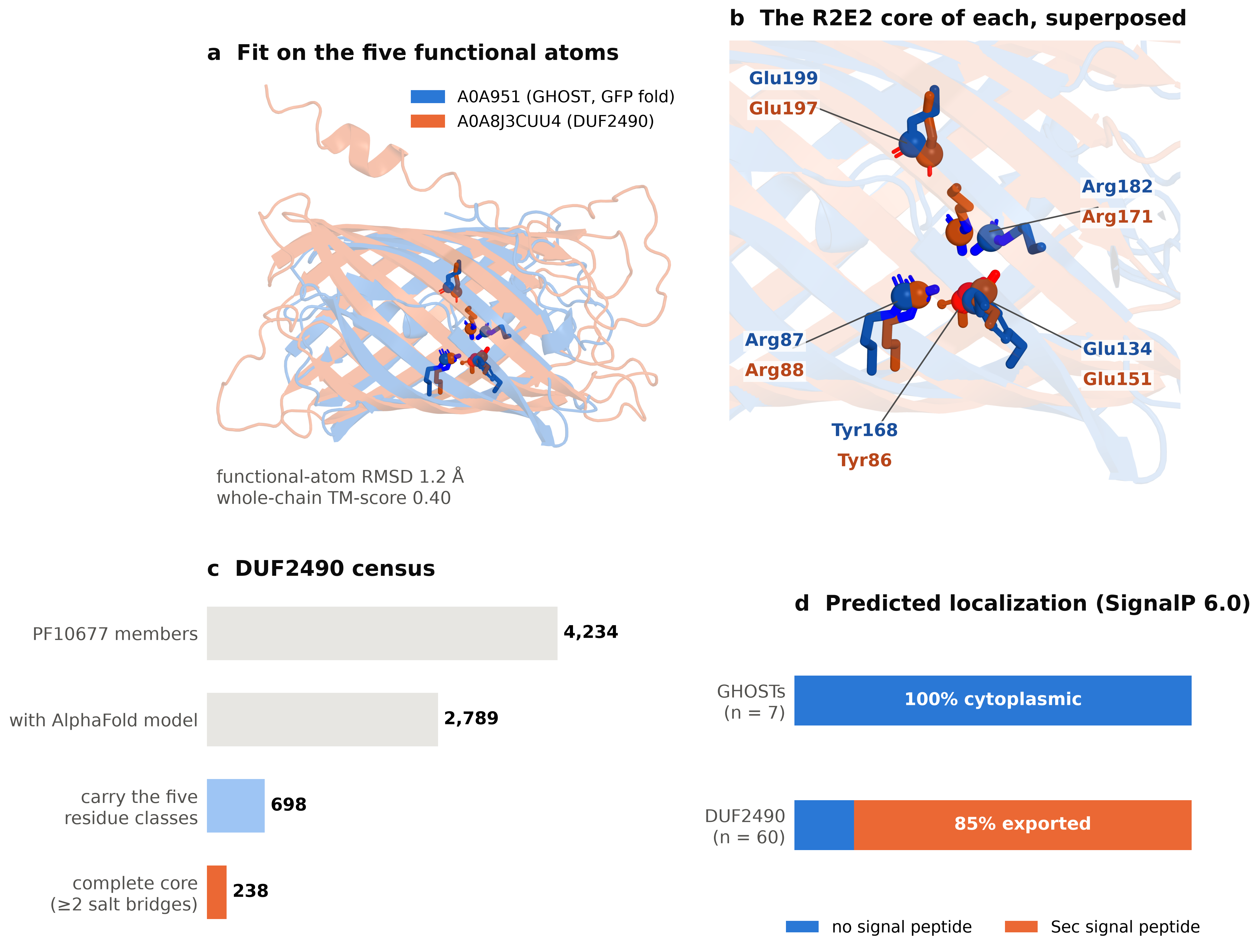

Supplementary Fig. 3 | Convergent recurrence of the buried R2E2 core on two unrelated β-barrel folds. (a) The A0A951 GFP-fold barrel (blue; GHOST founder, model AF-A0A951HUW2-F1) and a DUF2490 outer-membrane β-barrel (orange; UniProt A0A8J3CUU4, Pfam PF10677, AlphaFold model), superposed by least-squares fit of the five role-matched functional atoms only (Arg CZ, Glu CD, Tyr OH), not by global superposition. The functional atoms coincide with an RMSD of 1.2 Å, whereas the scaffolds diverge (whole-chain TM-score approximately 0.37-0.40, from a separate all-versus-all TM-align calculation): the compact GFP β-can with its central helix against the larger DUF2490 barrel with extended loops and an N-terminal helix. (b) Close-up of the superposed cores, with side chains as sticks and functional atoms as spheres. Each label pairs the A0A951 residue (blue) with its DUF2490 counterpart (orange): Arg87/Arg88, Glu134/Glu151, Tyr168/Tyr86, Arg182/Arg171 and Glu199/Glu197. (c) PF10677 census: of 4,234 UniProt members, 2,789 have AlphaFold models, 698 carry the two-cation, two-anion, aromatic constellation, and 238 meet the complete-core criterion of at least two salt bridges with the aromatic engaged below 4.0 Å (Supplementary Note 7). (d) SignalP 6.0 predictions: none of seven tested GHOST sequences carries a signal peptide, whereas 51 of 60 sampled DUF2490 sequences (85%) carry a Sec/SPI signal peptide, consistent with export to the outer membrane. The same buried core is therefore embedded in two structurally unrelated barrels sharing approximately 5-7% sequence identity, consistent with convergent evolution rather than common ancestry.

##### **S7.6 Predicted compartment split**

SignalP 6.0^48^ predicted no signal peptide for the seven tested GHOST proteins (OTHER probability approximately 1.0, including A0A951) and a Sec/SPI signal peptide for 51 of 60 sampled DUF2490 proteins (probabilities 0.95 to 0.98; predicted cleavage at residues 20 to 27). Together with the fold classification, these results predict cytoplasmic localization for the tested GHOST proteins and export of DUF2490 to the periplasm/outer membrane. The same buried network can therefore be maintained in contrasting bulk redox compartments, consistent with a compartment-insulated internal core.

##### **S7.7 Shared and distinct features of the two carriers**

Both carriers contain the R2E2 core in a beta-barrel lumen, a glutamine near one anion, a second aromatic near the central tyrosine, and substantial representation in Bacteroidota. They differ in detectable sequence homology, taxonomic breadth, predicted cellular compartment, barrel size, and scaffold architecture: the GHOST family is a soluble eleven-stranded GFP beta-can with a central helix, whereas DUF2490 is a larger predicted transmembrane outer-membrane barrel. Consistent with these differences, all-versus-all structural clustering places the GHOST family with canonical fluorescent proteins rather than with DUF2490. These contrasts identify the five-residue network, rather than the surrounding fold, as the recurring structural unit.

##### **S7.8 Caveats**

No experimental DUF2490 structure is available, so the census depends on AlphaFold side-chain placements, although the buried residues are modeled at high confidence. The AFDB50 FoldDisco index represents cluster representatives rather than all AlphaFold models; the complete PF10677 census covers every member with a retrievable model, but 1,445 sequences lacked models and were not screened. The structural and sequence evidence favors convergence but does not formally exclude very deep homology. Finally, the biochemical function of the network remains undemonstrated in both families.

#### **Supplementary Note 8 | Structural tolerance of substitutions at the chromophore-equivalent position**

##### **S8.1 Prediction design**

Residue 61 varies across the GHOST family while the buried charged/redox core remains strongly constrained. We therefore tested whether representative substitutions at this position altered the predicted A0A951 fold or pocket architecture. Six constructs were analyzed: WT A0A951HUW2 (M61), M61V, M61F, M61Y, M61T, and M61H. The panel includes naturally observed family-like hydrophobic or polar states and two mechanistic controls. M61Y and M61H are not treated as naturally observed family states on the basis of this analysis.

##### **S8.2 Barrel closure, confidence, and pocket packing**

All models retained the predicted barrel seam. Across every construct and pass, the mean number of inter-strand C-alpha contacts below 8 Å was 28 with zero between-model standard deviation. Construct/pass mean minimum seam distances ranged from 4.494 to 4.515 Å, and the full model-level range was 4.481-4.532 Å. Construct/pass mean whole-model pLDDT ranged from 92.912 to 93.505. Mean pLDDT for residues 58-90 ranged from 92.323 to 94.017, that for residues 178-202 ranged from 95.902 to 96.585, and mean cross-region PAE ranged from 2.956 to 3.455 Å.

No model had a residue-61-to-Tyr83 minimum heavy-atom distance below 2.4 Å. Short residue-61-to-Arg182 contacts occurred only in the template-free pass: M61V in 3 of 8 models, M61H in 2 of 8, M61T in 2 of 8, and M61F in 1 of 8. No such contact occurred for WT or M61Y, and none occurred for any construct in the GFP-template pass. The grid calculation found no large, reproducible pocket opening. Mean enclosed volumes were 0-2.25 Å³ across construct/pass means, and several groups had no detected enclosed component. Because components below 3 Å³ were excluded on a 1 Å grid and occasional individual models contained components up to 15 Å³, these values are comparative packing estimates rather than physical cavity measurements.

##### **S8.3 Compatibility ranking and template sensitivity**

WT ranked first under the heuristic compatibility score in both passes. The ordering of the variants changed modestly: without templates, M61H > M61F > M61V > M61Y > M61T; with templates, M61V > M61H > M61F > M61T > M61Y. The rank instability among variants, together with the uniformly small positive template shift of 0.51-1.17 score units, argues against a strong substitution-specific structural effect. The robust findings are preservation of barrel closure and high local confidence across all constructs, not the exact ordering of closely spaced scores.

M61Y was geometrically accommodated in both passes. It retained high regional confidence and low cross-region PAE, showed no short contact to Tyr83 or Arg182, and produced only very small enclosed-volume estimates. These calculations do not support gross steric incompatibility or loss of the predicted GFP-fold architecture as an explanation for failure of an M61Y construct to express or mature. They do not establish chromophore formation, fluorescence, folding in cells, or biochemical activity.

##### **S8.4 Interpretation limits**

Prediction seeds sample model behavior and are not experimental or biological replicates. All models were unrelaxed, and close side-chain contacts may change after local minimization. The template pass is a sensitivity analysis, not independent evidence for folding. AlphaFold and ColabFold do not simulate chromophore maturation, proton transfer, redox chemistry, or explicit internal solvent, and similar pLDDT values do not establish equal thermodynamic stability or expression yield. The compatibility score is a transparent comparative screen, not a free energy. Finally, the panel establishes tolerance in the A0A951 background only. Scored on the independent Hartley^83^ central-helix criterion, M61Y and M61F remain within the chromophore-forming range (0.650 and 0.654 A against a 1.0487 A threshold), but the equivalent tyrosine restoration in three other family scaffolds falls outside it: A0A4R7JUD5 T62Y 1.400 A, A0A495W682 T32Y 1.514 A and A0A931S450 T93Y 1.614 A. With all four parents re-predicted under the designs' own ColabFold protocol, three wild-type barrels lie clearly below the threshold (0.619, 0.844 and 0.910 A) and the fourth, A0A931S450, sits 0.012 A above it at 1.061 A, inside the 0.091 A control standard deviation and so not separable from the cut. All three tyrosine designs lie beyond it. The substitution therefore accounts for the separation outright in A0A495W682 and A0A4R7JUD5, and for most but not demonstrably all of it in A0A931S450. The panel's conclusion should therefore not be generalized across the family.

#### **Supplementary Note 9 | Environmental sources of GHOST sequences**

The 22 full-length members occur in oxic or sub-oxic, largely surface or photic, oxidatively exposed contexts. Seventeen are from free-living environmental microbes, two are host-encoded coral proteins, and three are host- or symbiont-associated members from sponge, macroalga, and lichen contexts. Nine occur in surface soil, four in terrestrial fungi, three in coral reefs, two in sub-oxic groundwater, two in other marine settings, one in a thermal spring, and one in the lichen *Lepraria neglecta*. No member comes from an obligately anoxic source or an anoxic host-internal setting such as the gut or digestive tract. Seven of 22 members (32%) are from explicitly extreme or high-irradiance niches: three coral-reef members, two polar-soil members, one thermal-spring member, and the desiccation/UV-tolerant lichen. The family spans all seven continents and both polar regions. Phenylalanine occupies the chromophore-equivalent site in all three coral-reef members, but is not confined to them: it is also carried by the iron-rich hot-spring member and by one of the groundwater members, giving five phenylalanine members across three habitats. The remaining members carry methionine, valine or threonine. The chromophore-position residue therefore does not track biome. These patterns are reported descriptively rather than as a formal enrichment test because available ecosystem tallies do not match the structure/profile discovery pipeline and the 22 sequences are phylogenetically non-independent.

Sample metadata was retrievable for 22 of the environmental sequences recovered by the exhaustive search, a set distinct from the 22 complete barrels of the analysed roster. Twelve of these 22 have known coordinates. Seven come from marine samples. All seven were identified in Tara Oceans protist size-fraction metatranscriptomes collected across four ocean basins between 36° N and 9° S. Because these are metatranscriptomic data, the genes were actively transcribed by eukaryotic plankton rather than simply being present as environmental DNA. Five members come from grassland or forest soils in California and Panama. One member was recovered independently from the same field site in Hopland, California, by two separate sequencing studies. Other members with identifiable sources extend the family into freshwater or estuarine sediment, a poplar root endosphere, and an algal wastewater bioreactor. These habitats had not previously been associated with the family.

#### **Supplementary Table 1 | The GHOST roster**

The 25 records analysed in this work: 22 complete barrels and three partial fragments (HEY8628922, MEY2932110, RKT55308). "Tyr66 site" is the residue at the position equivalent to avGFP Tyr66. Habitat follows the NCBI BioSample ecosystem field. A0A146F2F0 is excluded as a redundant gene model: A0A7R7WR58 is identical to it over all 254 residues. MBV9931914 is the same protein as the founder and is not listed separately. Entries deleted from UniProtKB are cited by UniParc identifier.

| **Accession** | **UniParc** | **Organism** | **Kingdom** | **Completeness** | **Length (aa)** | **Tyr66 site** | **Habitat** | **UniProt status** |
| --- | --- | --- | --- | --- | --- | --- | --- | --- |
| A0A4R7JUD5 | UPI00105F07CE | *Maribacter spongiicola* | Bacteria | complete barrel | 240 | T | marine | live (TrEMBL) |
| A0A931S450 | UPI001DD0F21D | Deltaproteobacteria bacterium | Bacteria | complete barrel | 261 | T | groundwater | DELETED from UniProtKB |
| A0A951HUW2 | UPI001DADDAB1 | Alphaproteobacteria bacterium | Bacteria | complete barrel | 218 | M | rhizosphere soil | DELETED from UniProtKB |
| GAB4285421 |  | Opitutales bacterium | Bacteria | complete barrel | 244 | F | hot spring | not checked |
| KGS53704 |  | *Burkholderia pseudomallei* MSHR5609 | Bacteria | complete barrel | 253 | V | environmental | not checked |
| MDB5144524 |  | *Mucilaginibacter* sp. | Bacteria | complete barrel | 237 | M | soil | not checked |
| MDP1614463 |  | Methylococcales bacterium | Bacteria | complete barrel | 245 | F | groundwater | not checked |
| MDQ5822267 |  | Actinomycetota bacterium | Bacteria | complete barrel | 215 | M | soil | not checked |
| MDX6514066 |  | Gaiellaceae bacterium | Bacteria | complete barrel | 214 | T | soil | not checked |
| MEP6978467 |  | Thermoleophilia bacterium | Bacteria | complete barrel | 215 | M | soil (polar) | not checked |
| MES2389882 |  | Bacteroidota bacterium | Bacteria | complete barrel | 241 | M | soil | not checked |
| WP_299115178 |  | uncultured *Winogradskyella* sp. | Bacteria | complete barrel | 223 | M | marine | not checked |
| XWB57685 |  | *Actinophytocola* sp. CA-135451 | Bacteria | complete barrel | 231 | T | soil | not checked |
| XWJ43021 |  | *Saccharothrix australiensis* | Bacteria | complete barrel | 231 | T | soil | not checked |
| A0A1M3T1J2 | UPI000924AB72 | *Aspergillus luchuensis* | Fungi | complete barrel | 253 | V | terrestrial fungal | DELETED from UniProtKB |
| A0A3F3PKF2 | UPI000E110040 | *Aspergillus welwitschiae* | Fungi | complete barrel | 253 | V | terrestrial fungal | live (TrEMBL) |
| A0A7R7WR58 | UPI001AFCCAD2 | *Aspergillus kawachii* | Fungi | complete barrel | 254 | V | terrestrial fungal | live (TrEMBL) |
| A0AAE0DQW0 | UPI00295C7812 | *Lepraria neglecta* | Fungi | complete barrel | 251 | V | terrestrial (lichen) | live (TrEMBL) |
| A0ABR1SEL9 | UPI00312D2A0A | *Apiospora marii* | Fungi | complete barrel | 263 | V | not recorded in master table | live (TrEMBL) |
| A0A7U3V473 | UPI001AF9A9FC | *Acropora tenuis* (coral) | Cnidaria | complete barrel | 216 | F | marine | live (TrEMBL) |
| XP_067048447 |  | *Acropora muricata* (coral) | Cnidaria | complete barrel | 216 | F | marine | not checked |
| MGYP000217628526 |  | environmental metagenome | environmental metagenome | complete barrel | 200 | F | marine (coral reef) | not checked |
| HEY8628922 |  | bacterial (fragment) | Bacteria | partial fragment | 154 | M | not recorded in master table | not checked |
| MEY2932110 |  | bacterial (fragment) | Bacteria | partial fragment | 159 | T | not recorded in master table | not checked |
| RKT55308 |  | bacterial (fragment) | Bacteria | partial fragment | 201 | T | not recorded in master table | not checked |

#### **Supplementary Table 2 | Cross-domain provenance and evidence strength**

Origin evidence for the eukaryotic and reef-associated members. Intron-containing genes and a cloned transcript strongly exclude bacterial reads mis-binned into eukaryotic assemblies; single-exon gene models and metagenomic records are interpreted more cautiously. Exon structures were read from the GenBank coding-sequence records named in the table.

| Member | Organism (lineage) | Source record | Source type | Exons | Protein existence | Evidence against bacterial contamination |
| --- | --- | --- | --- | --- | --- | --- |
| A0A7U3V473 | *Acropora tenuis* (Cnidaria, Scleractinia) | EMBL LC519770; BBV24626.1 | cloned mRNA transcript | n/a (mRNA) | PE 2, transcript level | Strong: cloned eukaryotic transcript |
| XP_067048447 | *Acropora muricata* (Cnidaria) | RefSeq XM_067192346.1; GeneID 136923871 | RefSeq genome-annotated gene | 1 (CDS 126..776) | RefSeq model mRNA | Moderate: annotated coral gene, no intron confirmation |
| A0A7R7WR58 | *Aspergillus kawachii* (Fungi) | EMBL AP024426; BCR95063.1; UP000661280 | reference-genome gene | 2 (join 7534..7777 / 7858..8378) | PE 3, homology | Strong: spliced gene architecture |
| A0A3F3PKF2 | *Aspergillus welwitschiae* (Fungi) | EMBL KZ852097; RDH27192.1; UP000253729 | reference-genome gene | 2 (complement join 48216..48736 / 48817..49057) | PE 3, homology | Strong: spliced gene architecture |
| A0AAE0DQW0 | *Lepraria neglecta* (Fungi, lichen-forming) | EMBL JASNWA010000004; KAK3176910.1; UP001276659 | reference-genome gene | 2 (join 3958428..3958668 / 3958727..3959241) | PE 3, homology | Strong: spliced gene architecture |
| A0ABR1SEL9 | *Apiospora marii* (Fungi) | EMBL JAQQWI010000006; KAK8032773.1; UP001396898 | reference-genome gene | 1 (68538..69329) | PE 3, homology | Moderate: annotated fungal gene, no intron confirmation |
| A0A1M3T1J2 | *Aspergillus luchuensis* (Fungi) | genome assembly; UniProtKB entry withdrawn (UniParc UPI000924AB72) | reference-genome gene | not confirmed | PE 3 | Moderate: record withdrawn, intron structure unconfirmed |
| MGYP000217628526 | environmental metagenome, coral-reef biome (MGnify ERP108071) | MGnify protein record | metagenomic | unknown | none | Source organism undetermined; reef-associated, not a coral gene |

The strongest cross-domain evidence is the cloned *Acropora tenuis* transcript and the intron-containing *Aspergillus kawachii*, *Aspergillus welwitschiae*, and *Lepraria neglecta* genes. The *Acropora muricata* record rests on a single-exon RefSeq gene model, the two *Aspergillus luchuensis* records require confirmation of gene structure, and MGYP000217628526 remains a coral-reef-associated metagenomic sequence whose source organism is unknown.

#### **Supplementary Table 3 | Custom-script reproducibility map**

Mapping of the custom repository scripts to the analyses described in the Methods. Paths are relative to the top-level public archive. Software versions, inputs, parameters, and principal outputs are retained with the repository records.

| Script (archive path scripts/) | Analysis implemented | Execution host |
| --- | --- | --- |
| 69_site_selection.py | Per-site purifying-selection analysis of the family | workstation |
| 74_md_setup_explicit.py | Build the explicit-solvent MD system for A0A951 (linear chromophore site) | workstation |
| 75_md_run_explicit.py | Run explicit-solvent production MD on a GPU | workstation |
| 77_run_water_md_gpu.sh | Driver: explicit-solvent water-occupancy MD | GPU workstation |
| 85_md_setup_immature.py | Parameterise the immature-ring residue CIM and build the solvated system | workstation |
| 86_run_immature_md_gpu.sh | Driver: immature-ring water-occupancy MD | GPU workstation |
| 105_md_setup_mature_acropora.py | Parameterise the mature CRO chromophore and build the coral F67Y system | workstation |
| 106_run_mature_acropora_md_gpu.sh | Driver: mature-chromophore coral F67Y MD | GPU workstation |
| 109_fig_sequence_logo.py | Two-group sequence logo at the diagnostic fingerprint positions (Fig. 2) | workstation |
| 110_strucmotif_pdb_csm_search.py | Fold-independent 3D motif search for the R2E2 core against the PDB and RCSB CSM | workstation |
| 111_folddisco_afdb50_motif.py | Fold-independent motif search of AFDB50 for the R2E2 core (FoldDisco) | workstation |
| 112_duf2490_census_screen.py | Full PF10677 (DUF2490) census screened for the R2E2 core | workstation |
| 113_fold_classification.py | Deep-homology versus convergence test for DUF2490 network carriers | workstation |
| 113_site_rates_mltree.py | Site-rate analysis on the ML tree (LG+G4) rather than the NJ tree | workstation |
| 114_run_mltree_iqtree_gpu.sh | Driver: IQ-TREE version of the ML-tree site-rate rerun | GPU workstation |
| 114_signalp_localization.py | SignalP 6.0 localization of both network-carrying families | workstation |
| 115_charge_network_map.py | Geometric map of the buried charge network, aromatic chain and metal check | workstation |
| 116_family_propka_signature.py | Conservation of the buried-Arg / bridging-Glu electrostatic signature | workstation |
| 117_stage_cavity_docking_gpu.sh | Driver: cavity, electrostatics and ligand-docking tier | GPU workstation |
| 118_environmental_ecology.py | Ecological analysis of the family | workstation |
| 119_gold_enrichment_test.py | Illustrative comparison of the ecological pattern to a GOLD background | workstation |
| 120_network_conservation.py | Frequency of the buried network across the 22 complete members | workstation |
| 121_core_vs_barrel_rate_test.py | Prespecified five-residue core tested against the rest of the barrel | workstation |
| 123_md_network_occupancy.py | Network contact occupancy over MD trajectories (Supplementary Table 5) | workstation |
| 124_run_md_network_occupancy_gpu.sh | Driver: MD network-occupancy analysis | GPU workstation |
| 133_render_signal_overlay_pub.py | Publication render of the cross-fold convergence overlay (Supplementary Fig. 3) | workstation |
| paper1_figures.py | Render the taxon-targeted ML phylogeny and the per-site evolutionary-rate figure | workstation |
| render_fig1_gpu.sh | Driver: render the barrel figure as two panels with headless PyMOL | GPU workstation |
| render_fig1a.py | Whole-protein overview of the A0A951 GFP-fold barrel | workstation |
| render_fig1b.py | Active-site close-up: chromophore-equivalent site and catalytic residues | workstation |
| render_fig4.pml | Per-site rate mapped as B-factor on the A0A951 barrel | GPU workstation |
| render_fig4_gpu.sh | Driver: render the conservation-mapped barrel | GPU workstation |
| render_paper1_figs_gpu.sh | Convenience wrapper for both structural figure renders | GPU workstation |
| render_signal_overlay.py | Cross-fold convergence overlay of the GFP-fold barrel and a DUF2490 barrel | workstation |
| render_signal_overlay_gpu.sh | Driver: render the cross-fold convergence overlay | GPU workstation |

#### **Supplementary Table 4 | Matched local-control tests for defense and mobile-element adjacency**

Focal GHOST loci were compared with length-matched, non-marker control proteins from the same retrievable genomic windows. Tests are one-sided in the enrichment direction; the paired permutation result is reported for mobile-element proximity. Twelve GHOST loci had retrievable local windows. Up to 20 controls were drawn per locus, matched within 0.70 to 1.30 times the focal protein length where possible, giving 103 controls in total; adjacency was scored over a window of plus or minus 10 genes, and control proteins that were themselves defense or mobile-element markers were excluded. No test is significant after accounting for the three features examined.

| Feature (+/-10 genes) | GHOST loci | Matched controls | Odds ratio | One-sided p | Paired permutation p |
| --- | --- | --- | --- | --- | --- |
| Defense-system adjacent | 2/12 (0.17) | 28/103 (0.27) | 0.63 | 0.875 | 0.916 |
| Mobile-element adjacent | 2/12 (0.17) | 7/103 (0.07) | 3.06 | 0.237 | 0.065 |
| Either defense or mobile adjacent | 4/12 (0.33) | 35/103 (0.34) | 1.02 | 0.633 | 0.436 |

The controls address local genomic context only. Source-habitat patterns were coded separately and remain descriptive because same-window controls share habitat metadata and the discovery pipeline lacks a matched database-wide denominator.

#### **Supplementary Table 5 | Molecular-dynamics contact occupancies**

Contact occupancies summarize three independent 100-ns replicas per system. Values for the two founder protonation states or two coral force-field variants are reported as the observed range where the source analysis combined them; a single value means the two agreed to within 0.005. Interactions are named in A0A951 model numbering. The equivalent coral A0A7U3V473 residues are Arg188, Glu140, Arg93, Tyr174, Glu205, Gln203, Ser172 and Tyr89. Distances are means over the pooled replicas of the first-listed condition.

| Interaction (A0A951 numbering) | Type | Founder immature ring, occupancy | Coral F67Y mature, occupancy | Founder distance (A) | Coral distance (A) |
| --- | --- | --- | --- | --- | --- |
| Arg182-Glu134 | salt | 1.00 | 1.00 | 2.71 +/- 0.07 | 2.72 +/- 0.08 |
| Arg87-Glu134 | salt | 1.00 | 0.47-0.48 | 2.78 +/- 0.12 | 3.94 +/- 1.01 |
| Arg182-chromophore-site backbone | hbond | 1.00 | 0.00 | 2.84 +/- 0.10 | 7.03 +/- 0.67 |
| Arg87-Tyr168 | hbond | 0.78-0.80 | 0.69-0.70 | 3.31 +/- 0.24 | 3.45 +/- 0.63 |
| Glu199-Gln197 | hbond | 0.97-1.00 | 0.00 | 2.91 +/- 0.48 | 7.98 +/- 1.45 |
| Arg87-Ser166 | hbond | 1.00 | 0.88-0.99 | 2.93 +/- 0.11 | 3.14 +/- 0.35 |
| Tyr83-Tyr168 | aromatic | 0.59-0.67 | 0.16-0.31 | 6.92 +/- 0.36 | 7.26 +/- 0.51 |

#### **Supplementary Table 6 | Residue-61 panel confidence and barrel-closure summaries**

Values are means across eight ColabFold seeds. Pocket pLDDT is shown for residues 58-90 and 178-202. Seeds sample prediction behavior and are not biological replicates.

| Template pass | Construct | Seeds | Mean pLDDT | Pocket pLDDT 58-90 | Pocket pLDDT 178-202 | Cross-region PAE | Barrel-end min CA (A) | Barrel-end contacts <8 A |
| --- | --- | --- | --- | --- | --- | --- | --- | --- |
| GFP templates | M61F | 8 | 93.2 | 93.1 | 96.2 | 3.22 | 4.51 | 28 |
| GFP templates | M61H | 8 | 93.2 | 93.3 | 96.3 | 3.21 | 4.50 | 28 |
| GFP templates | M61T | 8 | 93.2 | 92.9 | 96.1 | 3.31 | 4.50 | 28 |
| GFP templates | M61V | 8 | 93.3 | 93.2 | 96.3 | 3.18 | 4.50 | 28 |
| GFP templates | M61Y | 8 | 93.0 | 92.7 | 96.1 | 3.37 | 4.51 | 28 |
| GFP templates | WT | 8 | 93.5 | 94.0 | 96.6 | 2.96 | 4.49 | 28 |
| template-free | M61F | 8 | 93.1 | 92.9 | 96.1 | 3.32 | 4.51 | 28 |
| template-free | M61H | 8 | 93.1 | 93.0 | 96.2 | 3.29 | 4.51 | 28 |
| template-free | M61T | 8 | 93.0 | 92.3 | 95.9 | 3.46 | 4.51 | 28 |
| template-free | M61V | 8 | 93.1 | 92.7 | 96.0 | 3.34 | 4.51 | 28 |
| template-free | M61Y | 8 | 92.9 | 92.4 | 95.9 | 3.45 | 4.51 | 28 |
| template-free | WT | 8 | 93.4 | 93.6 | 96.4 | 3.10 | 4.50 | 28 |

#### **Supplementary Table 7 | Residue-61 pocket contacts, cavity estimates, and compatibility**

Values are means across eight ColabFold seeds. Short R182 is the percentage of models with a minimum residue-61-to-Arg182 heavy-atom distance below 2.4 Å. No residue-61-to-Tyr83 short contacts occurred. Cavity is the mean total enclosed grid volume. Delta is the GFP-template compatibility score minus the corresponding template-free value and is therefore defined only for the GFP-template rows. Short Arg182 contacts occur only in the template-free pass and never for the wild type or M61Y.

| Template pass | Construct | 61-83 (A) | 61-87 (A) | 61-182 (A) | 61-197 (A) | 61-199 (A) | Short R182 (%) | Cavity total (A3) | Compatibility | Delta vs template-free |
| --- | --- | --- | --- | --- | --- | --- | --- | --- | --- | --- |
| GFP templates | M61F | 9.34 | 4.08 | 3.72 | 5.29 | 4.08 | 0 | 0.38 | 188.5 | +0.6 |
| GFP templates | M61H | 9.22 | 3.95 | 3.83 | 5.40 | 4.07 | 0 | 1.75 | 188.7 | +0.5 |
| GFP templates | M61T | 9.38 | 3.61 | 3.46 | 5.27 | 4.06 | 0 | 2.25 | 187.9 | +1.0 |
| GFP templates | M61V | 9.39 | 3.46 | 2.93 | 5.33 | 3.96 | 0 | 0.00 | 188.8 | +1.2 |
| GFP templates | M61Y | 9.68 | 3.26 | 4.35 | 5.21 | 4.03 | 0 | 0.88 | 187.6 | +0.7 |
| GFP templates | WT | 9.42 | 3.98 | 5.17 | 5.35 | 3.73 | 0 | 0.50 | 190.3 | +0.9 |
| template-free | M61F | 9.63 | 3.82 | 3.55 | 5.25 | 4.09 | 12 | 1.38 | 187.9 | +0.6 |
| template-free | M61H | 9.28 | 3.88 | 2.85 | 5.43 | 4.12 | 25 | 0.00 | 188.2 | +0.5 |
| template-free | M61T | 9.42 | 3.14 | 2.86 | 5.69 | 4.17 | 25 | 0.38 | 186.9 | +1.0 |
| template-free | M61V | 9.45 | 3.30 | 2.46 | 5.44 | 4.04 | 38 | 0.00 | 187.6 | +1.2 |
| template-free | M61Y | 9.86 | 3.11 | 3.25 | 5.25 | 4.10 | 0 | 1.25 | 186.9 | +0.7 |
| template-free | WT | 9.50 | 4.01 | 4.38 | 5.41 | 3.81 | 0 | 0.00 | 189.3 | +0.9 |

#### **Supplementary Data files | Principal repository outputs**

Paths are relative to the root of the archived data set (Data availability). Folders 01_query_and_models/ to 10_figures/ hold the processed data for the main analyses. 11_motif_search_convergence/ holds the motif-search and DUF2490 outputs: census/DUF2490_all_members_uniprot.tsv, census/DUF2490_full_network_screen.tsv, census/DUF2490_complete_core_members.tsv, hit_sets/ (RCSB and FoldDisco results), relatedness_TM_seqid.md, localization/ (SignalP predictions) and models/ (representative AlphaFold models). 12_residue61_panel/ holds the residue-61 ColabFold panel: inputs/panel.fasta, scripts/run_panel.sh, runs/manifest.tsv, the three run configurations (runs/msa/, runs/no_templates/ and runs/gfp_templates/), analysis/per_model.tsv, analysis/per_construct.tsv, analysis/report.md, scripts/analyze_panel.py and docs/Colab.md. scripts/ holds the analysis and figure-generation scripts mapped in Supplementary Table 3.

#### **Supplementary Table 8 | Additional GHOST candidates recovered beyond the 25-record analysed roster**

Members recovered beyond the originally characterized set. Eighteen full-length metagenomic members (sequence side) carry the complete eight-residue buried-network fingerprint; one further member was found on the structure side. TM is to the *Acropora* query. Every chromophore-equivalent seat is non-tyrosine. The table lists 20 rows: the 18 sequence-side members with the complete fingerprint, the single structure-side member with the complete fingerprint (A0A6V0IBN8), and A0AAN0NCD3, which is listed for completeness at 6/8 and is not counted as a family member.

| Member | Source | TM to query | Chromophore seat (avGFP Y66) | Network score | Status |
| --- | --- | --- | --- | --- | --- |
| APDOM4702015248_1054824 | JGI scaffold | 0.86 | V | 8/8/8 | sequence side, complete fingerprint |
| ERR1712176_929985 | SRA metagenome | 0.79 | V | 8/8/8 | sequence side, complete fingerprint |
| ERR1719189_151448 | SRA metagenome | 0.79 | F | 8/8/8 | sequence side, complete fingerprint |
| ERR1719193_33278 | SRA metagenome | 0.86 | V | 8/8/8 | sequence side, complete fingerprint |
| ERR1719277_550285 | SRA metagenome | 0.85 | V | 8/8/8 | sequence side, complete fingerprint |
| ERR1719343_925520 | SRA metagenome | 0.86 | V | 8/8/8 | sequence side, complete fingerprint |
| ERR1719356_1157905 | SRA metagenome | 0.79 | V | 8/8/8 | sequence side, complete fingerprint |
| ERR1719507_167900 | SRA metagenome | 0.81 | F | 8/8/8 | sequence side, complete fingerprint |
| EndMetStandDraft_8_1072994 | JGI scaffold | 0.85 | V | 8/8/8 | sequence side, complete fingerprint |
| LauGreSBDMM110SN_4 | JGI scaffold | 0.86 | V | 8/8/8 | sequence side, complete fingerprint |
| MGYP000547889416 | MGnify | 0.99 | F | 8/8/8 | sequence side, complete fingerprint |
| SRR4051812_44845058 | SRA metagenome | 0.96 | M | 8/8/8 | sequence side, complete fingerprint |
| SRR6266540_7379509 | SRA metagenome | 0.93 | M | 8/8/8 | sequence side, complete fingerprint |
| UPI000536FBBC | UniParc | 0.79 | V | 8/8/8 | sequence side, complete fingerprint |
| UPI0005381B41 | UniParc | 0.79 | V | 8/8/8 | sequence side, complete fingerprint |
| UPI0018F23B0B | UniParc | 0.92 | M | 8/8/8 | sequence side, complete fingerprint |
| UPI00232C93DA | UniParc | 0.81 | V | 8/8/8 | sequence side, complete fingerprint |
| tagenome__1003787_1003787 | JGI scaffold | 0.86 | V | 8/8/8 | sequence side, complete fingerprint |
| A0A6V0IBN8 | AlphaFold DB (UniParc) | 0.88 | (model not local) | 8/8 | structure side, complete fingerprint |
| A0AAN0NCD3 | AlphaFold DB | ~ | (model not local) | 6/8 | structure side, incomplete fingerprint (not counted as a member) |

#### **Supplementary Table 9 | Per-structure R2E2 core geometry**

Functional-atom distances between R2E2 core positions and relative solvent accessibility (RSA) of the five core residues, for 16 GHOST models and 12 canonical fluorescent proteins carrying all five core residues (matched controls), measured on predicted structures with positions transferred from A0A951 by pairwise alignment. Medians, GHOST against control: R182-E134 4.40 against 16.67 Å; R182-E199 7.65 against 11.34 Å; mean RSA 0.01 against 0.11. These values underlie Fig. 4.

| **Structure** | **Set** | **R87-E134 (Å)** | **R182-E134 (Å)** | **R182-E199 (Å)** | **R87-Y168 (Å)** | **Y168-E199 (Å)** | **mean RSA** | **max RSA** |
| --- | --- | --- | --- | --- | --- | --- | --- | --- |
| A0A4R7JUD5 | GHOST | 5.66 | 4.19 | 6.70 | 3.98 | 10.07 | 0.006 | 0.025 |
| A0A7U3V473 | GHOST | 5.86 | 5.88 | 7.11 | 4.44 | 11.02 | 0.009 | 0.031 |
| A0A931S450 | GHOST | 5.06 | 4.55 | 8.25 | 3.43 | 10.89 | 0.011 | 0.051 |
| A0A951HUW2 | GHOST | 5.79 | 3.93 | 7.93 | 4.26 | 10.98 | 0.004 | 0.011 |
| GAB4285421 | GHOST | 5.95 | 4.25 | 9.10 | 4.38 | 12.16 | 0.004 | 0.014 |
| KGS53704 | GHOST | 6.18 | 3.58 | 8.37 | 4.31 | 11.46 | 0.015 | 0.041 |
| MDB5144524 | GHOST | 5.87 | 5.68 | 6.73 | 4.78 | 10.52 | 0.001 | 0.004 |
| MDP1614463 | GHOST | 4.97 | 5.89 | 6.87 | 3.98 | 11.00 | 0.002 | 0.010 |
| MDQ5822267 | GHOST | 6.67 | 3.80 | 8.23 | 5.20 | 10.93 | 0.002 | 0.005 |
| MDX6514066 | GHOST | 5.55 | 3.80 | 8.03 | 3.96 | 10.59 | 0.011 | 0.041 |
| MEP6978467 | GHOST | 6.40 | 4.08 | 7.80 | 5.10 | 10.68 | 0.005 | 0.025 |
| MES2389882 | GHOST | 5.75 | 6.01 | 6.94 | 4.40 | 10.57 | 0.000 | 0.000 |
| RKT55308 | GHOST | 5.73 | 5.18 | 7.50 | 3.77 | 10.41 | 0.009 | 0.035 |
| WP_299115178 | GHOST | 5.93 | 5.59 | 6.72 | 4.46 | 10.84 | 0.003 | 0.014 |
| XP_067048447 | GHOST | 6.00 | 5.85 | 6.76 | 4.53 | 10.83 | 0.010 | 0.026 |
| XWJ43021 | GHOST | 5.92 | 3.90 | 7.82 | 3.88 | 10.41 | 0.015 | 0.045 |
| A0A182DW87 | matched control | 6.87 | 16.53 | 11.32 | 4.25 | 9.92 | 0.103 | 0.455 |
| A0A9Y2YAB8 | matched control | 6.99 | 17.96 | 11.53 | 4.70 | 10.11 | 0.134 | 0.618 |
| A0AAN0NCD3 | matched control | 7.11 | 16.59 | 11.36 | 4.76 | 9.81 | 0.107 | 0.487 |
| A0AAU9WCI9 | matched control | 6.30 | 18.08 | 11.74 | 4.45 | 9.19 | 0.122 | 0.571 |
| A8CLT7 | matched control | 6.01 | 17.99 | 12.04 | 4.17 | 9.26 | 0.137 | 0.614 |
| B9UPG6 | matched control | 6.24 | 17.24 | 11.77 | 4.21 | 8.94 | 0.102 | 0.464 |
| D0VX33 | matched control | 6.77 | 16.66 | 11.33 | 4.54 | 10.20 | 0.111 | 0.500 |
| H3JQU4 | matched control | 6.81 | 16.69 | 11.21 | 4.65 | 9.94 | 0.114 | 0.521 |
| H3JQU7 | matched control | 7.00 | 16.63 | 11.02 | 4.84 | 9.87 | 0.120 | 0.536 |
| Q7Z168 | matched control | 6.19 | 17.82 | 11.68 | 4.22 | 8.88 | 0.112 | 0.506 |
| Q963I9 | matched control | 6.40 | 16.18 | 11.15 | 4.30 | 9.49 | 0.119 | 0.576 |
| Q9U6Y6 | matched control | 5.62 | 13.69 | 11.02 | 4.23 | 9.55 | 0.096 | 0.444 |

#### **Supplementary Table 10 | Per-structure Poisson-Boltzmann potentials**

Electrostatic potential at the chromophore-site backbone carbonyl oxygen from APBS (PARSE charges, pH 7.0), with the charges of the chromophore-forming triad zeroed, for 17 GHOST models, 18 general fluorescent proteins and 12 matched core carriers. Means: GHOST 36.2, general 19.4 and matched 24.8 kT/e. The Arg-zeroed potential was computed for the GHOST and general sets only (n.d., not determined). These values underlie the electrostatics section of the Results. This 47-structure PARSE calculation is separate from the single-founder AMBER calculation described in the Supplementary Methods.

| **Structure** | **Set** | **Tyr66-site residue** | **φ (kT/e)** | **φ, Arg side chains zeroed (kT/e)** | **nearest Arg (Å)** |
| --- | --- | --- | --- | --- | --- |
| A0A4R7JUD5 | GHOST | THR | 39.8 | -41.4 | 2.65 |
| A0A7U3V473 | GHOST | PHE | 13.0 | -42.8 | 3.29 |
| A0A931S450 | GHOST | THR | 34.3 | -43.1 | 3.22 |
| A0A951HUW2 | GHOST | MET | 46.6 | -35.3 | 2.88 |
| GAB4285421 | GHOST | PHE | 59.5 | -31.1 | 2.46 |
| KGS53704 | GHOST | VAL | 44.3 | -22.7 | 3.20 |
| MDB5144524 | GHOST | MET | 36.5 | -31.0 | 3.28 |
| MDP1614463 | GHOST | PHE | 34.6 | -36.7 | 3.05 |
| MDQ5822267 | GHOST | MET | 31.2 | -35.9 | 3.50 |
| MDX6514066 | GHOST | THR | 32.2 | -35.2 | 3.25 |
| MEP6978467 | GHOST | MET | 34.0 | -37.0 | 3.31 |
| MES2389882 | GHOST | MET | 41.6 | -24.1 | 3.35 |
| RKT55308 | GHOST | THR | 30.4 | -35.8 | 3.60 |
| WP_299115178 | GHOST | MET | 19.2 | -46.3 | 3.46 |
| XP_067048447 | GHOST | PHE | 36.8 | -36.4 | 2.87 |
| XWB57685 | GHOST | THR | 43.5 | -38.7 | 2.62 |
| XWJ43021 | GHOST | THR | 37.0 | -40.3 | 2.79 |
| B1PNC0 | general fluorescent protein | TYR | 19.3 | -33.6 | 3.12 |
| D0VWW2 | general fluorescent protein | TYR | 22.1 | -1.7 | 4.20 |
| D1MPT3 | general fluorescent protein | TYR | 11.0 | -23.9 | 4.17 |
| P42212 | general fluorescent protein | TYR | 31.4 | 7.4 | 4.20 |
| P83690 | general fluorescent protein | TYR | 21.3 | -22.4 | 4.05 |
| Q5S6Z9 | general fluorescent protein | TYR | 29.4 | -27.8 | 3.57 |
| Q5TLG6 | general fluorescent protein | TYR | 36.4 | -24.9 | 3.45 |
| Q8I6J8 | general fluorescent protein | TYR | 25.3 | -31.5 | 3.50 |
| Q8ISF8 | general fluorescent protein | TYR | 8.6 | -10.0 | 4.64 |
| Q95P04 | general fluorescent protein | TYR | 16.1 | -26.1 | 4.08 |
| Q95W85 | general fluorescent protein | TYR | 9.1 | -29.4 | 4.22 |
| Q95W86 | general fluorescent protein | TYR | -0.3 | -22.5 | 4.20 |
| Q9GZ28 | general fluorescent protein | TYR | -0.4 | -25.1 | 4.16 |
| Q9U6Y4 | general fluorescent protein | TYR | 2.3 | -60.7 | 3.19 |
| Q9U6Y5 | general fluorescent protein | TYR | 18.7 | -32.6 | 3.29 |
| Q9U6Y6 | general fluorescent protein | TYR | 26.1 | -34.3 | 3.07 |
| Q9U6Y7 | general fluorescent protein | TYR | 16.1 | -7.0 | 4.18 |
| Q9U6Y8 | general fluorescent protein | TYR | 57.5 | 36.6 | 4.56 |
| A0A182DW87 | matched core carrier | n.d. | 51.0 | n.d. | n.d. |
| A0A9Y2YAB8 | matched core carrier | n.d. | 33.0 | n.d. | n.d. |
| A0AAN0NCD3 | matched core carrier | n.d. | 35.9 | n.d. | n.d. |
| A0AAU9WCI9 | matched core carrier | n.d. | 34.8 | n.d. | n.d. |
| A8CLT7 | matched core carrier | n.d. | 52.4 | n.d. | n.d. |
| B9UPG6 | matched core carrier | n.d. | 28.8 | n.d. | n.d. |
| D0VX33 | matched core carrier | n.d. | -20.6 | n.d. | n.d. |
| H3JQU4 | matched core carrier | n.d. | 11.3 | n.d. | n.d. |
| H3JQU7 | matched core carrier | n.d. | 8.3 | n.d. | n.d. |
| Q7Z168 | matched core carrier | n.d. | 29.7 | n.d. | n.d. |
| Q963I9 | matched core carrier | n.d. | 16.1 | n.d. | n.d. |
| Q9U6Y6 | matched core carrier | n.d. | 16.6 | n.d. | n.d. |

#### **Supplementary Table 11 | Sequence identity of each complete GHOST barrel to avGFP**

Identities from a BLOSUM62 global pairwise alignment to avGFP (PDB 1EMA, 238 residues; gap-open -10, gap-extension -0.5). The four columns differ only in denominator; the paper uses aligned residue pairs throughout. Rows are sorted by that column.

| Member | Length (aa) | Identical residues | % / aligned pairs | % / shorter sequence | % / avGFP length | % / alignment columns |
| --- | --- | --- | --- | --- | --- | --- |
| A0A951HUW2 | 218 | 44 | 22.8 | 20.2 | 18.5 | 16.7 |
| MES2389882 | 241 | 53 | 24.9 | 22.3 | 22.3 | 19.9 |
| A0A931S450 | 261 | 53 | 25.2 | 22.3 | 22.3 | 18.3 |
| A0ABR1SEL9 | 263 | 53 | 25.6 | 22.3 | 22.3 | 18.0 |
| A0A7R7WR58 | 254 | 50 | 25.8 | 21.0 | 21.0 | 16.8 |
| A0A3F3PKF2 | 253 | 51 | 26.0 | 21.4 | 21.4 | 17.3 |
| XWJ43021 | 231 | 47 | 26.0 | 20.3 | 19.7 | 16.3 |
| WP_299115178 | 223 | 57 | 26.8 | 25.6 | 23.9 | 23.0 |
| A0AAE0DQW0 | 251 | 55 | 28.5 | 23.1 | 23.1 | 18.6 |
| A0A4R7JUD5 | 240 | 57 | 29.4 | 23.9 | 23.9 | 20.1 |
| KGS53704 | 253 | 53 | 29.4 | 22.3 | 22.3 | 17.0 |
| MGYP000217628526 | 200 | 53 | 29.9 | 26.5 | 22.3 | 20.3 |
| MDB5144524 | 237 | 60 | 30.0 | 25.3 | 25.2 | 21.8 |
| XWB57685 | 231 | 54 | 30.0 | 23.4 | 22.7 | 18.7 |
| A0A1M3T1J2 | 253 | 60 | 30.5 | 25.2 | 25.2 | 20.4 |
| GAB4285421 | 244 | 63 | 30.9 | 26.5 | 26.5 | 22.7 |
| A0A7U3V473 | 216 | 58 | 31.0 | 26.9 | 24.4 | 21.7 |
| XP_067048447 | 216 | 58 | 31.4 | 26.9 | 24.4 | 21.6 |
| MEP6978467 | 215 | 60 | 31.7 | 27.9 | 25.2 | 22.7 |
| MDQ5822267 | 215 | 57 | 32.6 | 26.5 | 23.9 | 20.5 |
| MDX6514066 | 214 | 59 | 33.9 | 27.6 | 24.8 | 21.2 |
| MDP1614463 | 245 | 60 | 34.5 | 25.2 | 25.2 | 19.4 |

### **Data availability**

Data and code availability. The canonical family roster, with live primary accessions, UniParc identifiers for the three entries no longer in UniProtKB (A0A951HUW2, A0A1M3T1J2 and A0A931S450), taxon, completeness, fingerprint state and the analyses each record enters, is provided as Supplementary Table 1. Per-structure measurements behind Fig. 4 (functional-atom distances and relative solvent accessibility for 16 family and 12 identity-matched control models) and the per-structure Poisson-Boltzmann potentials for both control sets are provided as Supplementary Tables 9 and 10. The 54-sequence alignment, the maximum-likelihood tree with SH-aLRT and ultrafast bootstrap support, and the constraint trees and approximately unbiased test outputs are deposited with the manuscript. AlphaFold models for control proteins were retrieved from the AlphaFold Protein Structure Database (version 6). All processed data, analysis scripts and the residue-61 panel are archived at Zenodo [ADD DOI] and mirrored on GitHub [ADD URL]; paths below and in the Supplementary Information are relative to the root of that archive.

Query and models: 1EMA; AlphaFold A0A951HUW2. Source data: BioProject PRJNA687273, MAG BioSample SAMN17140656, source-metagenome BioSample SAMN10290267, SRA run SRR8113284, contig JAFAZW010000077.1, protein MBV9931914. Genomic-context and defence-system outputs are in 09_genomic_context_defense/ (A0A951HUW2_gene_neighborhood.md, A0A951HUW2_crossmember_neighborhoods.md, A0A951HUW2_defense_systems.md and A0A951HUW2_padloc_systems.md). The family profile HMM (a0a_subfamily.hmm), its seed alignment (a0a_subfamily_seed.afa), calibration, the UniProtKB and MGnify sweep and verification of cnidarian, fungal and metagenomic hits are in 03_sequence_homology_hmm/. Structural models and the per-member master table (01_query_and_models/), structure searches, phylogenies and ancestral reconstructions, site rates, core geometry, molecular-dynamics occupancy tables (07_molecular_dynamics/), electrostatics and figures occupy folders 01 to 10. Motif searches, the DUF2490 census, fold-classification tests and SignalP outputs are in 11_motif_search_convergence/. The residue-61 mutational panel, with exact sequences, run configurations, manifests, all 96 model-level measurements, construct summaries and the analysis code, is in 12_residue61_panel/. Analysis and figure scripts are in scripts/, mapped to analyses in Supplementary Table 3. Quantum-chemical analyses are detailed in the companion study.
